# Generation of excitatory and inhibitory neurons from common progenitors via Notch signaling in the cerebellum

**DOI:** 10.1101/2020.03.18.997205

**Authors:** Tingting Zhang, Tengyuan Liu, Natalia Mora, Justine Guegan, Mathilde Bertrand, Ximena Contreras, Andi H. Hansen, Carmen Streicher, Marica Anderle, Natalia Danda, Luca Tiberi, Simon Hippenmeyer, Bassem A. Hassan

## Abstract

Brain neurons arise from relatively few progenitors capable of giving rise to an enormous diversity of neuronal types. Nonetheless, a cardinal feature of mammalian brain neurogenesis in both the cortex and the cerebellum is that excitatory neurons and inhibitory neurons derive from separate, spatially segregated, progenitors. Whether bi-potential progenitors with an intrinsic capacity to generate both excitatory and inhibitory lineages exist and how such a fate decision may be regulated is unknown. Using cerebellar development as a model, we discover that individual embryonic cerebellar progenitors give rise to both inhibitory and excitatory lineages. We find that gradations of Notch activity levels determine the fates of the progenitors and their daughters. Daughters with the highest levels of Notch activity retain the progenitor fate. Daughters with intermediate levels of Notch activity become fate restricted to generate inhibitory neurons, while daughters with very low levels of Notch signaling adopt the excitatory fate. Therefore, Notch mediated binary cell fate choice is a mechanism for regulating the ratio of excitatory to inhibitory neurons from common progenitors.

**Graphical summary:** 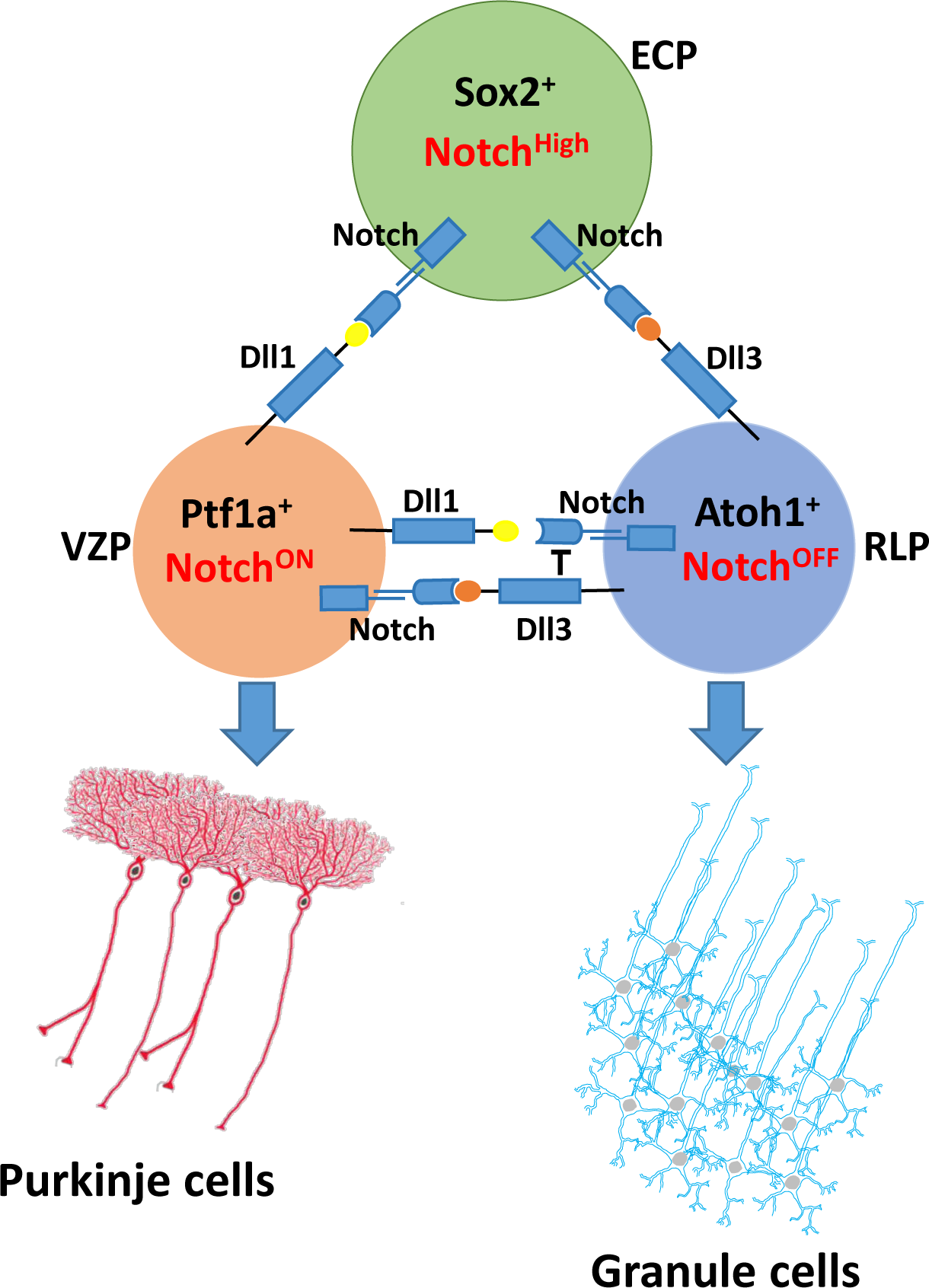

## INTRODUCTION

Correct brain function depends on neuronal diversity, whereby neurons of different morphologies and physiologies connect to produce functional neuronal circuits. In turn, neuronal diversity is generated from a relatively small and temporally limited number of bi-potential progenitor cells, or developmental neural stem cells. A major challenge in neurobiology is to understand how bi-potential progenitors endow their daughters of with different cell fates. The study of neurogenesis in a variety of models of vertebrates like zebrafish and mouse, as well as invertebrates like C. elegans and Drosophila, has converged on three major processes as key to understanding neuronal diversification. First, in both systems temporal mechanisms ensure that progenitors change competence over time to give rise to different type of neurons as brain development proceeds, as has been shown originally in the fly embryonic ventral nerve cord (Cleary and Doe, 2006; Doe, 2017) and subsequently in the fly visual system (Erclik et al., 2017; Li et al., 2013b) and mammalian neocortex (Oberst et al., 2019; Telley et al., 2016). Second, the combination of spatially restricted expression of cell fate determinants and temporal competence windows creates neuronal diversity by confining the generation of specific subtypes of neurons to specific neurogenic zones at different times. Elegant examples of how such combinatorial diversification have recently been reported for both the fly visual system and the mammalian neocortex (Apitz and Salecker, 2018; Erclik et al., 2017; Mayer et al., 2018; Mi et al., 2018; Mora et al., 2018; Nowakowski et al., 2017; Quan et al., 2016). Third, in *Drosophila* ample evidence demonstrates that individual neural progenitors can give rise to differentially fate-restricted daughters while also maintaining their own progenitor status through a binary cell fate choice mechanism mediated by the highly conserved Notch signaling pathway (Endo et al., 2007; Wheeler et al., 2008). In both the mammalian neocortex (Bonnefont et al., 2019; Castro et al., 2006; Imayoshi et al., 2010) and the cerebellum (Machold et al., 2007) there is evidence that Notch activity is required to maintain progenitors in a bi-potential state by preventing their differentiation. In this scenario, Notch signaling contributes to the temporal axis of neuronal diversity by maintaining progenitors as they pass through consecutive fate competence windows. In contrast, there is currently no evidence that common progenitors use Notch signaling to generate neuronal diversity in the mammalian brain through a binary cell fate choice mechanism, leading to speculation that this aspect of neurogenesis is regulated differently in inverterbrates and mammals.

The cerebellum is a hub for control of motor function and contributes to a number of higher brain functions such as reward-related cognitive processes (Carta et al., 2019; Kostadinov et al., 2019; Wagner and Luo, 2020). Deficits in cerebellar development lead to severe neurological disorders such as cerebellar ataxias (Manto et al., 2020) and medulloblastomas (Northcott et al., 2019), a heterogeneous and severe groups of childhood brain tumors. Thus, it is important to understand the underlying cellular and molecular mechanisms controlling cerebellar development. Until recently, the consensus view of the development of the cerebellum (Leto et al., 2016) suggested that different types of cerebellar neurons arise at different time points from two spatially distinct progenitor pools located in a continuum along the fourth ventricle, either dorsally in the rhombic lip (RL) or ventrally in the ventricular zone (VZ). Two basic helix-loop-helix (bHLH) transcription factors called Atonal homologue 1 (Atoh1; RL) and Pancreas transcription factor 1 alpha (Ptf1a; VZ), mark these progenitor (Figure S1A) domains (du Plessis et al., 2017; Wang and Zoghbi, 2001). During embryonic development Atoh1^+^ RL precursors give rise to glutamatergic neurons, while Ptf1a^+^ VZ precursors give rise to GABAergic neurons (Leto et al., 2016).

Despite what appear to be independent spatial and temporal origins, a number of intriguing observations suggest that these precursors may share some common features. First, the cell fate in the two germinal niches can be switched when Atoh1 and Ptf1a are ectopically expressed in the VZ and RL, respectively (Wang et al., 2005; Yamada et al., 2014). Second, pseudo-time trajectory analysis of single-cell RNAseq (scRNAseq) embryonic mouse cerebellum data suggests that a common pool of progenitors branches into either a glutamatergic fate or a GABAergic fate (Vladoiu et al., 2019). Third, the classic neural stem cell marker Sox2 is known as an early VZ marker ( Pibiri et al., 2016a), cells expressing both Sox2 and the glutamatergic RL fate determinant Atoh1 have been observed in human cerebellar organoids (Muguruma et al., 2015), suggesting that Sox2^+^ progenitors exist in both germinal zones. Very recent fate mapping of Sox2^+^ progenitors in the RL has shown that they can give rise to excitatory neurons (Selvadurai et al., 2020).

The fact that Sox2^+^ cells can give rise to both excitatory and inhibitory lineages is not inconsistent with the current consensus view of spatial and temporal segregation of progenitor domains (Leto et al., 2016). However, the studies highlighted above can also be interpreted to suggest that at least a subset of these Sox2^+^ cells are in fact bi-potential embryonic cerebellar progenitors (ECPs) each of which can simultaneously generate both excitatory and inhibitory lineages. Whether such ECPs exist, and how a binary excitatory versus inhibitory fate decision is regulated, are unknown. Here we use sparse fate mapping and loss and gain of function of Notch approaches in the mouse cerebellum and human cerebellar organoids, and demonstrate that excitatory and inhibitory lineages can derive from common progenitors and that Notch activity is required to for this decision.

## RESULTS

### GABAergic and glutamatergic neurons are generated simultaneously from Sox2^+^ progenitor pool

To identify markers for potential ECPs we examined gene expression in the embryonic cerebellum in the Allen Brain Atlas for well-established neural progenitor markers known to be expressed at various stages of cerebellar development (Figure S1C), namely Sox2 (Ahlfeld et al., 2017; Kelberman et al., 2008; Pibiri et al., 2016; Selvadurai et al., 2020), Nestin (Andreotti et al., 2018; Li et al., 2013a; Wojcinski et al., 2017), GLAST (Bauer et al., 2012; Miyazaki et al., 2017; Yamada et al., 2000), S100β (Hachem et al., 2007; Landry et al., 1989) and GFAP (Vong et al., 2015; Wen et al., 2013; Yang et al., 2008). We found that both Sox2 and Nestin are highly expressed in both the VZ and RL during early cerebellar neurogenesis, although Nestin expression appeared to be more sparse, consistent with previous fate mapping work showing the Nestin expressing progenitors principally give rise to late born interneurons and glial cells (Fleming et al., 2013; Wojcinski et al., 2017). We retrieved and analyzed a single cell RNAseq dataset (ENA: PRJEB23051 data set) (Carter et al., 2018) and found ∼2000 cells expressing Sox2 of which only 50% express Nestin (Figure S1D and S1E). Together with its more abundant expression in the tissue, this suggests that Sox2 is the best candidate for finding bi-potential ECPs capable of giving rise to both excitatory and inhibitory lineages. We verified this with antibody staining and found the Sox2 protein is broadly expressed throughout the cerebellar anlagen (E9.5 – E16; Figure S1F-S1J). Furthermore, some Sox2^+^ cells express Atoh1, which is expressed in the cerebellar primordium as early as E9.5 (Figure S1F’-S1J’). When the progenitor cells in the RL begin to migrate to form EGL zone, the Sox2 expression became weaker and weaker over time, suggesting a dynamic process of cell differentiation (Figure S1F-S1J’).

The cerebellar cortex is composed of a well-defined limited number of cell types with characteristic locations and morphologies (Figure S1B and S2A). To begin testing the pluripotency of Sox2^+^ progenitors, we performed lineage tracing by using *Sox2^CreERT2^/Gt(ROSA)26Sor^tdTomato^/Atoh1^GFP^* pregnant mice injected with very low doses of Tamoxifen (TAM, 0.05mg) at early embryonic stages (E11.5) and examined the adult cerebellum at P21. Consistent with the presence of bi-potential progenitors, we recovered most known cerebellar cortex neuronal and glial cell types in all the known layers (Figure S2B-S2H’).

In the mammalian cerebellum, Purkinje cells (PCs) and Granule cells (GCs) are the major GABAergic and glutamatergic neurons (Hibi and Shimizu, 2012; Wang and Zoghbi, 2001). PCs are born during E10.5-E13, but don’t express the classical postmitotic marker Calbindin until E14.5 (Goldowitz et al., 1997; Morales and Hatten, 2006). While, Pax6 is a marker for GCs, which is highly expressed in the RL at E11.5, and its strong expression can be observed in both outer and inner EGL until the postnatal stages (Divya et al., 2016). Focusing on GCs and PCs, we repeated the lineage tracing experiments lowering the TAM dose to 0.03mg at E10.5, the lowest dose that we found to reliably induce recombination in our mice and examined the cerebellar cortex at P21 using Pax6 and Calbindin, respectively. Even at this very low dose, we recovered both GCs and PCs in our clones (Figure 1A-1D) as well as inhibitory interneurons labeled with Pax2 (Figure 1E-1G”). Furthermore, at E15 we find that the progeny of Sox2^+^ progenitors include all the known precursor cell types (Figure S3), as judged by their respective markers: Calbindin for nascent Purkinje cells (Figure S3A and S3B), Atoh1 and Pax6 (for excitatory precursors; Figure S3C and S3D), and Pax2 (for precursors of GABAergic interneurons; Figure S3E-S3G”). Glutamatergic deep cerebellar nuclei (DCNs) are produced in the RL at E10.5-12.5 and migrate rostrally in a subpial stream (SPS) to enter the nuclear transitory zone (NTZ) (Fink et al., 2006). Olig2 is a marker for all DCN projection neurons, while Tbr1 and Pax6 are specific markers for glu-DCN neurons (Fink et al., 2006; Ju et al., 2016). Immunostaining results showed that Olig2^+^, Pax6^+^ and Tbr1^+^ cells all co-localized with Tomato^+^ cells at E15 (Figure S3H-S3N).

**Figure 1.**
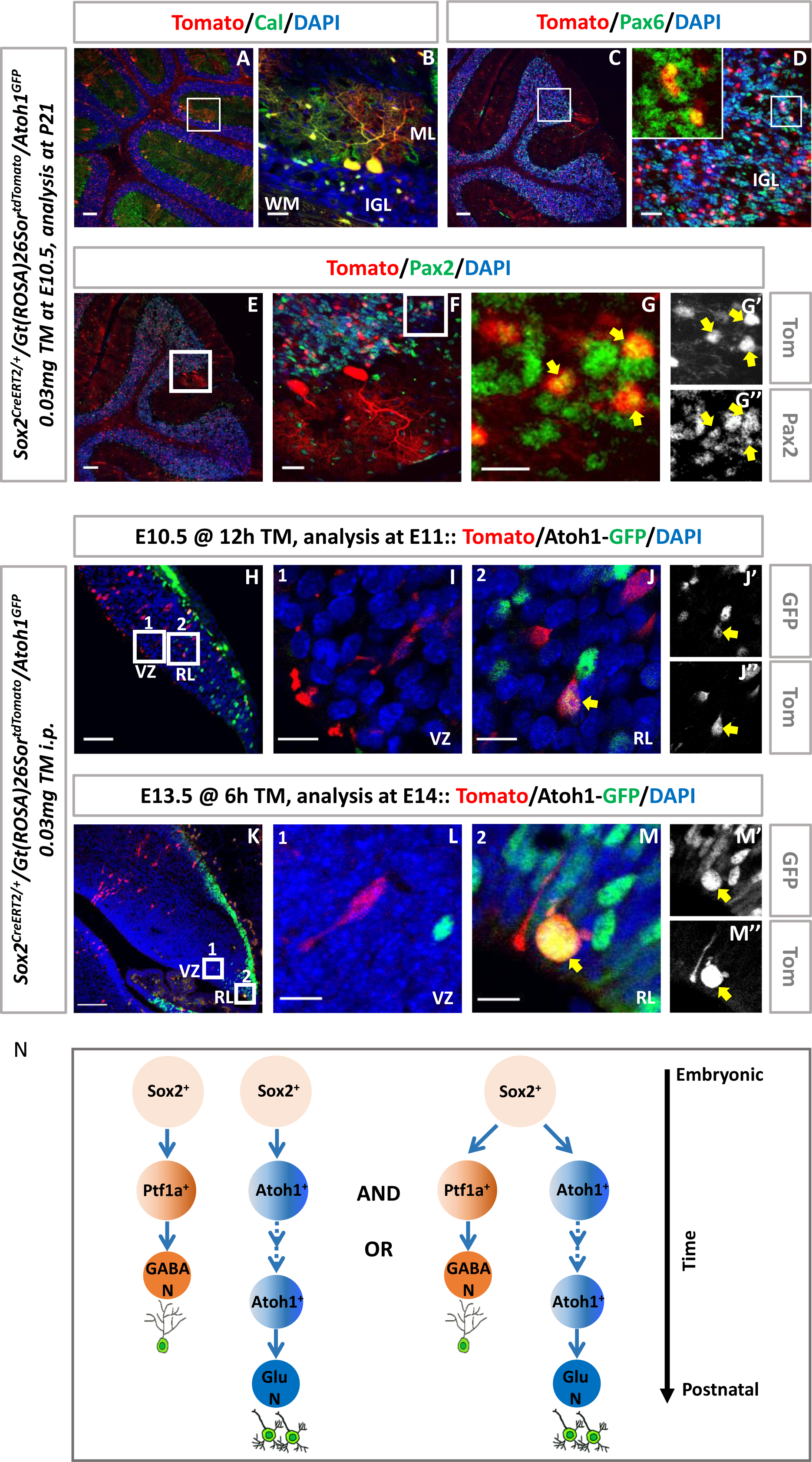
GABAergic and Glutamatergic lineages are generated simultaneously from Sox2^+^ progenitors: (A and C) Co-localization of tdTomato (red) with the PC marker Calbindin (Cal, green, A) or the GC marker Pax6 (green, C) in mouse cerebellum at P21. Scalebars=100 µm. (B and D) High magnification of the rectangular regions in (A) and (C). Scalebars= 25 µm. (E and F) Co-localization of the IN marker Pax2 (green) and tdTomato (red) in the mouse cerebellum at P21. Scalebars=100 µm and 25 µm. (G-G’’) Higher magnification of the rectangular region in (F). Scalebars= 25 µm. (H) Lineage tracing of Sox2^+^ cells (labeled by tdTomato, red) at E11 that 12h after injection of 0.03mg tamoxifen. Scalebars=50 µm. (I-J’’) High magnification of the rectangular regions in (H) which show the tdTomato expression in the VZ (I) and RL (J-J’’), respectively. Scalebars=10 µm. (K) Lineage tracing of Sox2^+^ cells (labeled by tdTomato, red) at E14 that 6h after injection of 0.03mg tamoxifen. Scalebars=100 µm. (L-M’’) High magnification of the rectangular regions in (K) which show the tdTomato expression in the VZ (L) and RL (M-M’’), respectively. Scalebars=15 µm. (N) Schematic of the two hypotheses that how Sox2^+^ progenitors generate both GABAergic and glutamatergic lineages in mouse cerebellum. Yellow arrows indicate Pax2^+^/ Tomato^+^ double-positive cells (G-G’’) or Sox2^+^/Atoh1-GFP^+^ double-positive cells (J-J’’ and M-M’’). Nuclei were stained with DAPI (blue).

The scRNAseq data suggested that relatively few Sox2^+^ cells persist as late as ∼E13. To increase the sparseness of our clones and examine them as soon as possible after the first cell division, we injected 0.03mg of TAM at both at E10.5 and E13.5 and examined embryos 6 hours and 12 hours later to ensure we focus on the immediate progeny of the labeled progenitors. As expected, with a low TAM dose and a short time induction, the combination resulted in rare labelled cells both at RL and VZ at E11 (Figure 1H-1J’’), and even fewer labelled cells at E14 (Figure 1K-1M’’). Surprisingly, however, these cells distributed equally between the EGL and the VZ (Figure 1K-1M’’), strongly suggesting that precursors of the two lineages are generated simultaneously. Although the nature of this lineages tracing approach cannot exclude the possibility that these very sparse clones derive from separate fate-restricted progenitors that divide simultaneously, the fact the we observe both PC and GC lineages within 6hrs of sparse labeling does suggest the existence of bi-potential progenitors even as late as E13.5 that divide and simultaneously give rise to both inhibitory and excitatory cells in the cerebellar primordium (Figure 1N).

### GABAergic and glutamatergic neurons are generated from single Sox2^+^ ECPs

To investigate whether such bi-potent progenitors exist, we asked if single Sox2^+^ ECPs can generate PCs and GCs *in vivo*. To this end, we turned to the clonal Mosaic Analysis with Double Markers (MADM) technology (Beattie et al., 2017; Gao et al., 2014; Zong et al., 2005). MADM allows the tracing of the progeny of single progenitors by virtue of the mode of segregation of two different markers (one green and one red) that cannot be expressed except following Cre-induced recombination during the G2 phase of the cell cycle.

If the two markers segregate to two different daughter cells, in what is called the G2-X event (Figure S4A), the progeny of the two daughters will be labelled in two different colors within the same brain. In our case, if G2-X recombination occurs in a progenitor that will give rise to 1 Atoh1^+^ daughter and 1 Ptf1a^+^ daughter and both survive, this will create clones containing both PCs and GCs labelled in distinct colors (for example green GCs and red PCs). If G2-X recombination occurs in a self-renewing ECP that itself will give rise to a bi-potential ECP which will then divide again to give rise to 1 Atoh1^+^ daughter and 1 Ptf1a^+^ daughter and both survive, this will create clones containing both PCs and GCs labelled in the same color (e.g. green GCs and green PCs). Any and all clones meeting the conditions described above are direct and conclusive evidence for the existence of a common progenitor for the two lineages. Conversely, clones that contain only one cell type suggest that recombination occurred in a fate-restricted progenitor.

Alternatively, if the two markers segregate together, in what is called the G2-Z event (Figure S4B), then one daughter and its progeny will be labelled in yellow and the other will not be labelled at all, and thus will not be detectable. In our case, if G2-Z recombination occurs in a self-renewing ECP that itself will give rise to a bi-potential ECP which will then divide again to give rise to 1 Atoh1^+^ daughter and 1 Ptf1a^+^ daughter and both survive, this will create clones containing both PCs and GCs labelled in yellow. Any yellow clones containing both PCs and GCs also provide conclusive direct evidence for a bi-potential progenitor for the two lineages. Single cell type yellow clones (for example yellow GCs and unlabeled PCs) are consistent with, but do not demonstrate the existence of, bi-potential progenitors. In summary, any and all clones that contain both cell types in yellow, both cell types in red, both cell types in green, or one cell type in red and the other in green, conclusively demonstrate a common cell of origin for GCs and PCs.

We performed MADM in cerebellar Sox2^+^ cells both at E10 and E11 and analyzed samples for GC and/or PC clones in adult mice. Consistent with sparsity of MADM clones (Beattie et al., 2017; Gao et al., 2014), we obtained 0.42 MADM events per brain hemisphere when injecting TM at E10 (32 clones in 76 brain hemispheres, Table 1), and 0.27 events when injecting TM one day later (23 clones in 86 brain hemispheres, Table 1). In 9 of the 55 clones (16.36%) we obtained both GCs and PCs in the same brains, with 6 clones being products of G2-X events and the remaining 3 products of G2-Z events with both PCs and GCs labelled in yellow (Table 1, Figure 2J-2L). Among G2-X clones, we obtained 2 scenarios: (i) G2-X recombination occurred in progenitors that generated two fate restricted daughter cells such that at P21 we obtained red labelled GCs and green labelled PCs in the same clone (Figure 2A-2C); and (ii) G2-X recombination occurred in self-renewing progenitors that generated one bi-potential ECP which then generated same color labelled PCs and GCs (i.e. either both in green or both in red), and another fate restricted Ptf1a^+^ cell that generated fate-restricted GABAergic lineage in the other color (Figure 2D-2I). These 9 clones that contained both PCs and GCs provide direct conclusive evidence that excitatory and inhibitory neurons can derive from a single bi-potent common progenitor.

**Figure 2.**
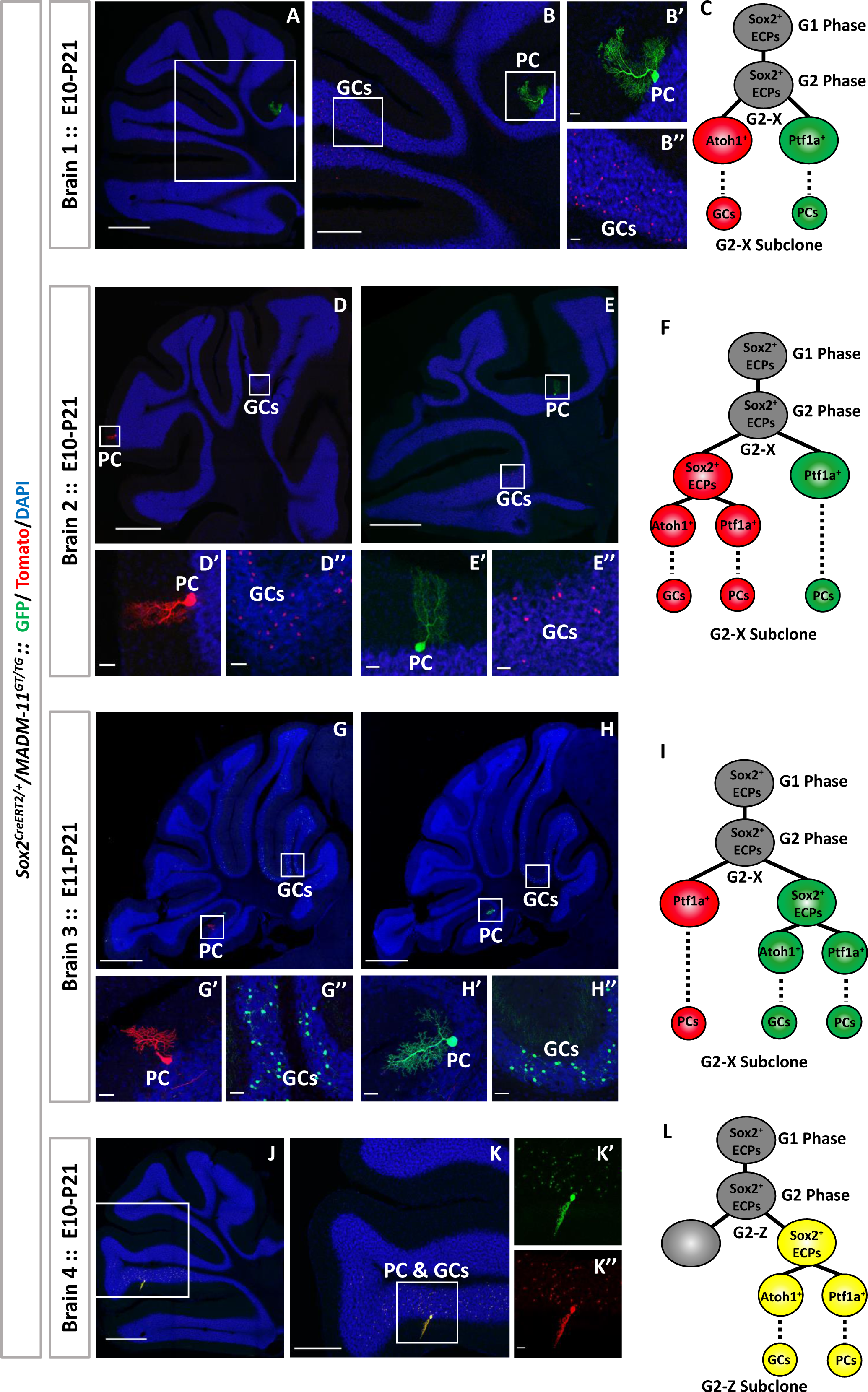
Sparse MADM-based lineage tracing of individual ECPs in mouse cerebellum: (A) Single progenitor G2-X-MADM clone at P21. One green PC and tens of red GCs were found in the same cerebellum. Scalebar=500 µm. (B) Higher magnification of the rectangular regions in (A). Scalebar=250 µm. (B’-B’’) Higher magnification of the rectangular regions in (B). Scalebars=25 µm (C) Scheme of a single G2-X-MADM-clone generated both fate restricted red (GCs) and green (PCs) lineages. (D and E) Single progenitor G2-X-MADM clone at P21. One red PC (D) and one green PC (E) and tens of red GCs were found in the same cerebellum. Scalebars=500 µm. (D’-E’’) Higher magnification of the rectangular regions in (D and E). Scalebars=25 µm. (F) Scheme of a single G2-X-MADM-clone generated a green fate restricted daughter cell (PCs lineage) and a red multipotent progenitor cell. (G and Single progenitor G2-X-MADM clone at P21. One red PC (G) and one green PC (H) and tens of green GCs were found in the same cerebellum. Scalebars=500 µm. (G’-H’’) Higher magnification of the rectangular regions in (G and H). Scalebars=25 µm. (I) Scheme of a single G2-X-MADM-clone generated a red fate restricted daughter cell (PCs lineage) and a green multipotent progenitor cell. (J) Single progenitor G2-Z-MADM clones at P21. One yellow PC and tens of yellow GCs were found in the same cerebellum. Scalebar=500 µm. (K) Higher magnification of the rectangular regions in (J). Scalebar=250 µm. (K’-K’’) Higher magnification of the rectangular regions in (K). Scalebars=25 µm. (L) Scheme of a single G2-Z-MADM-clone generated a yellow bi-potential progenitor cell and the other one without labeling. Nuclei were stained with DAPI (blue).

**Table 1.** MADM positive clones in mouse cerebellar hemisphere.

| E10-P21 | Number |  |  |  | Ratio |
| --- | --- | --- | --- | --- | --- |
| Total hemisphere | 76 |  |  |  |  |
| Total Clones | 32 |  |  |  | 42.10% |
| Clones contain PC and GC | 6 | Yellow | Green and Red | Green or Red | 18.75% |
|  |  | 3 | 2 | 1 |  |
| Clones contain PC only | 24 | Yellow | Green and Red | Green or Red | 75.00% |
|  |  | 8 | 6 | 10 |  |
| Clones contain GC only | 2 | Yellow | Green and Red | Green or Red | 6.25% |
|  |  | 2 | 0 | 0 |  |
| E11-P21 | Number |  |  |  | Ratio |
| Total hemisphere | 86 |  |  |  |  |
| Total Clones | 23 |  |  |  | 26.74% |
| Clones contain PC and GC | 3 | Yellow | Green and Red | Green or Red | 13.04% |
|  |  | 0 | 3 | 0 |  |
| Clones contain PC only | 11 | Yellow | Green and Red | Green or Red | 47.83% |
|  |  | 3 | 4 | 4 |  |
| Clones contain GC only | 9 | Yellow | Green and Red | Green or Red | 39.13% |
|  |  | 6 | 0 | 3 |  |

In addition, we obtained yellow clones containing either GCs only (8 clones, 14.55%) or PCs only (11 clones, 20%), consistent with a G2-Z event in a single progenitor cell (Figure S4K-S4L’, Table 1). In all of our single progenitor clones, we observed a much larger number of GCs than PCs, consistent with the transient amplification of GC precursors prior to neurogenesis (Espinosa and Luo, 2008). Finally, we also obtained (10/55, 18.18%) green and red single PCs cell type clones (Table1, Figure S4F-S4G) indicating recombination occurred in Sox2^+^ cells that had already committed to a specific Ptf1a^+^ lineage, and also (17/55, 30.91%) green or red single cell type clones (Table1, Figure S4H-S4J’) (Ahlfeld et al., 2017; Kelberman et al., 2008; Pibiri et al., 2016; Selvadurai et al., 2020).

### Single Sox2^+^ ECPs have an intrinsic potential to generate both PCs and GCs

To test if Sox2^+^ ECPs are intrinsically bi-potential, we carried out *in vitro* primary culture of cerebellar progenitors. We harvested ECPs from *Sox2^CreERT2^/Gt(ROSA)26Sor^tdTomato^* E11.5 embryos, sparsely labeled subsets of them by adding TAM to the culture, and performed time-lapse video microscopy for 12 consecutive days to trace the fate of individual Sox2^+^ ECPs (Figure 3A). Cells were kept under proliferative conditions for the first 3 days of culture to allow clonal expansion of single Sox2^+^ ECPs, followed by switching to a pro-differentiation medium for another 9 days to induce the production of both granule and Purkinje neurons (Kawasaki et al., 2000; Su et al., 2006). We traced the formation of 39 clones and found that 12 of them (31%) contained both PCs (Calbindin^+^/Tomato^+^) and GCs (Pax6^+^/Tomato^+^) derived from the same progenitor (Figure 3B-3G’’’ and Movie S1 and S2). These data demonstrate that a single Sox2^+^ ECP has an intrinsic ability to generate into both inhibitory and excitatory neurons.

**Figure 3.**
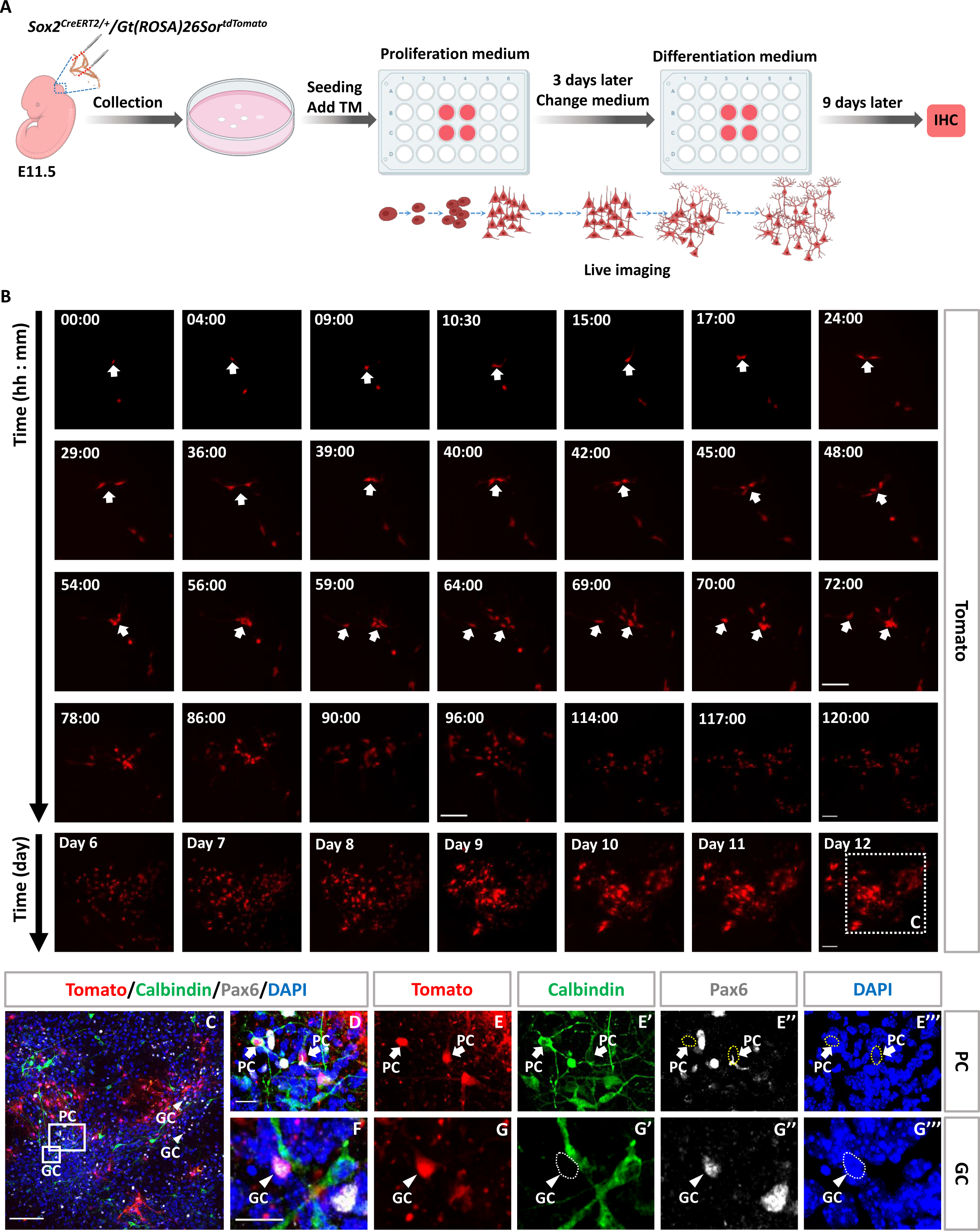
GABAergic and Glutamatergic lineages are generated from single Sox2^+^ progenitor in vitro: (A) Schematic representation of the mouse cerebellar progenitors primary culture for cell proliferation and differentiation. (B) Time-lapse video recording showed that a single Sox2 progenitor cell (Tomato^+^, Red) could divide several times to generate a cluster. Arrows indicate the Tomato^+^ cells that we focused on. (C) Immunolabeling of both PC marker (Calbindin, green) or GC marker (Pax6, gray) co-localized with Tomato (red) 9 days after cell differentiation, respectively. Arrow heads indicate Pax6^+^ / Tomato^+^ double positive cells. (D-G’’’) Higher magnification of the rectangular region in (C). Arrows indicate Calbindin^+^ / Tomato^+^ double positive cells and arrow heads indicate Pax6^+^ / Tomato^+^ double positive cells. Nuclei were stained with DAPI (blue). Scalebars=100 µm and 15 µm.

Finally, we asked if a common origin of GCs and PCs is conserved in human. We generated human-iPS-derived cerebellar organoid using a previously described protocol (Ishida et al., 2016; Muguruma et al., 2015). In particular the cerebellar organoids expressed midbrain-hindbrain markers such as EN*2* and GBX2 on day 21 (Muguruma et al., 2015). When iPS-derived cerebellar organoids were further cultured until day 25-26, they showed a few Sox2^+^ cells (Figure S5A) and Kirrel2^+^ cells that is among the earliest markers for the cerebellar neuroepithelium (Muguruma et al., 2015). Therefore, in order to sparsely label early Sox2^+^ human ECPs, we electroporated pGL3-Sox2Cre and pCAG PiggyBac (Pbase) and pPBCAG-LSL-Venus vectors into cerebellar organoids at 25 days (Figure S5A-S5B”’ and S5G). At day 35 the cerebellar organoids expressed several markers such as Ptf1a, Atoh1, Skor2, Pax6 (Ballabio et al., 2020a; Muguruma et al., 2015) but only from day 38-41 we were able to detect Calbindin^+^ cells. Therefore, we traced the Sox2^+^ ECPs fate at 41 days of organoid culture (Figure S5G). In all organoids, we observed both PCs (Venus^+^/Calbindin^+^, Figure S5C-S5D”’) and GCs (Venus^+^/Pax6^+^, Figure S5E-S5F’’’).

In summary, together our data thus far provide direct evidence for the existence of Sox2^+^ embryonic cerebellar progenitors capable of generating both GABAergic and Glutamatergic lineages.

### Differential expression of Notch pathway genes in different embryonic cerebellar progenitors

To identify the molecular underpinnings of the diversification of cerebellar neurons from common progenitors, we first analyzed available single-cell RNA sequencing (scRNAseq) data to define the molecular features of different cerebellar cell types during early stages. We retrieved the original data from E10-E13 (ENA: PRJEB23051 data set) (Carter et al., 2018), and used the Chromium system (10x Genomics) to profile the populations of each individual developmental time points. According to known cerebellar markers and transcriptional similarity, we identified individual clusters and mapped pseudo-time trajectory for cerebellar lineages (Figure 4A-4D). This confirmed that Sox2^+^ progenitors can give rise to two major lineages: glutamatergic cells from RL and GABAergic cells from VZ (Figure 4A). Previous work manipulating Notch1 function throughout the cerebellar anlagen identified Notch1 as required for the maintenance of RL progenitors by acting as an antagonist of RL-derived neurogenesis through the suppression of Atoh1 expression (Machold et al., 2007). We asked whether the role of Notch signaling in Sox2^+^ ECPs is only to contribute to the temporal axis of neurogenesis by maintaining progenitor fate, as is its classic role in mammalian neurogenesis, or whether it is also involved in the binary excitatory versus inhibitory cell fate decision. During canonical Notch-dependent binary cell fate decisions, cells interact to adopt mutually exclusive “opposing” terminal fates (Artavanis-Tsakonas et al., 1999a). We therefore began by testing whether the expression of Notch pathway genes correlates with cell fate in the scRNAseq data. We identified individual clusters for Notch signaling related genes, and found that canonical Notch target genes Hes5 and Hes1 are highly expressed in Ptf1a^+^ and Sox2^+^ cells, when compared with Atoh1^+^ cells, both at E12 and E13 (Figure 4E-4H). We next examined the expression of Notch1 and 5 of its ligands in these three populations (Table S1). We found that ECPs (green) express high levels of the Notch1, Hes1 and Hes5 mRNAs but very low levels of the ligands Dll1 and Dll3 (Figure 4I-4R). Ptf1a^+^ inhibitory precursors (red) express Notch1, Hes1 and Hes5 to a lesser extent than ECPs, but much higher levels of the trans-activating ligand Dll1 and slightly higher levels of the cis-inhibiting ligand Dll3 (Ladi et al., 2005) (Figure 4I-4R). Atoh1^+^ excitatory precursors (blue) express the lowest levels of the Notch1 receptor but the highest levels of the ligands, especially Dll3 (Figure 4I-4R). Taken together, analysis of Notch signaling related genes in these three populations revealed that Sox2^+^ ECPs have the highest level of Notch activity, followed by Ptf1a^+^ inhibitory precursors, with Atoh1^+^ excitatory precursors have the lowest levels of Notch activity, but the highest levels of Notch ligands. These observations suggest that Notch activity segregates the three populations from each other: first the bi-potential ECPs from their fate-restricted Ptf1a^+^ and Atoh1^+^ daughters and then Ptf1a^+^ inhibitory precursors from Atoh1^+^ excitatory precursors. We set out to test this idea by manipulating Notch activity specifically within Sox2^+^ ECPs.

**Figure 4.**
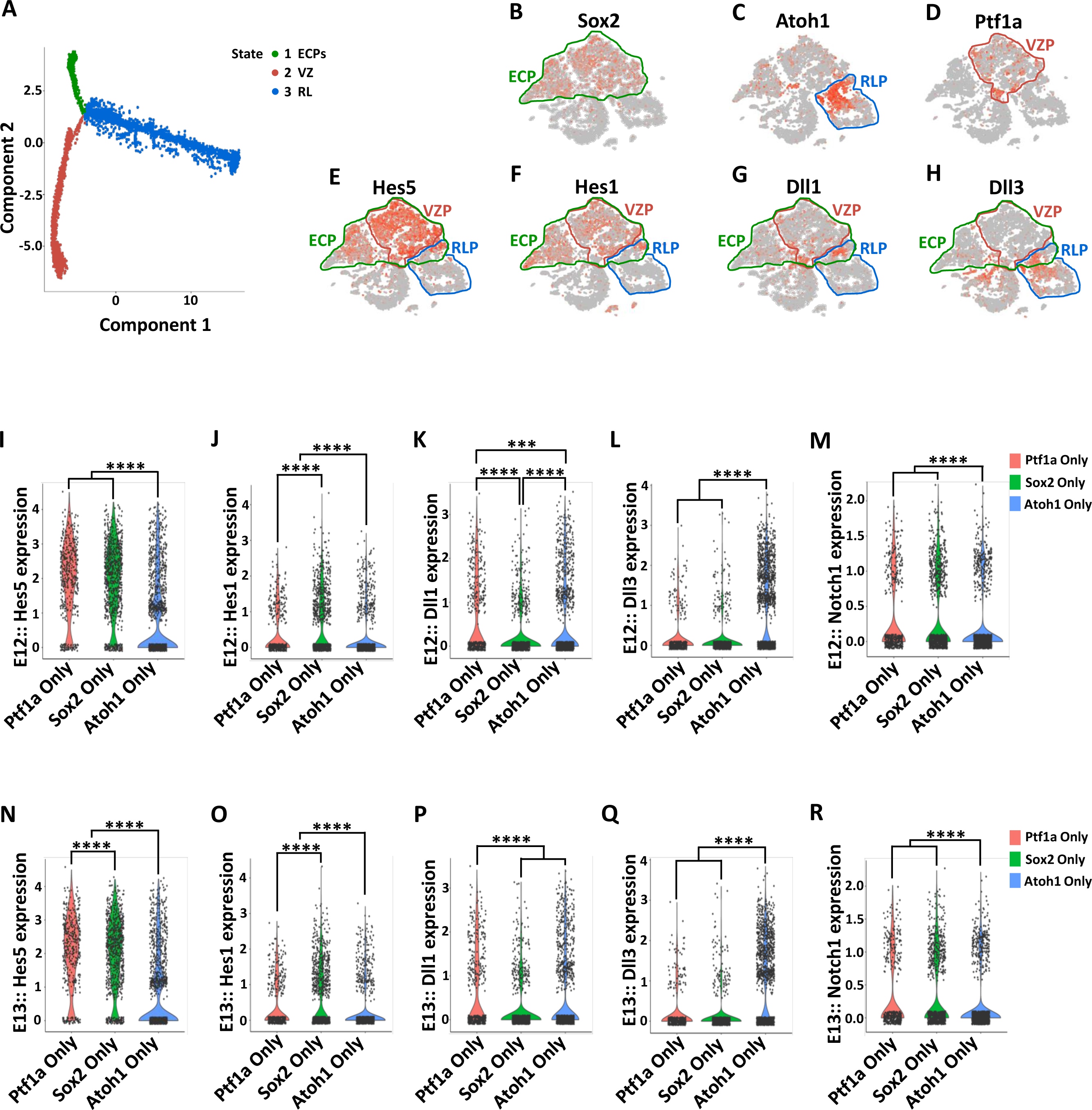
Analysis of Notch signaling gene expression in scRNAseq date from embryonic cerebellar progenitors: (A) Pseudo-time trajectory of scRNAseq data in the E10-E13 cerebellum: ECPs (green), VZ GABAergic lineages (red) and RL glutamatergic lineages (blue). (B) t-SNE visualizations of cerebellar derived cell clusters at E10-E13. Cells that express only Sox2 but neither Atoh1 nor Ptf1a are defined as ECPs (inside the green line). (C and D) t-SNE shows cell type specific marker expression: Atoh1 and Ptf1a. Atoh1^+^ clusters are rhombic lip progenitors (RLP, inside the blue line) and Ptf1a^+^ clusters are ventricular zone progenitors (VZP, inside the red line). Cells were color-coded according to gene expression with the cells expressing the gene indicated colored in orange-red. (E-H) t-SNE showing the expression of Notch signaling genes: Hes5, Hes1, Dll1 and Dll3. (I-R) Violin Plot showed the expression of Hes5, Hes1, Dll1, Dll3 and Notch1 in three different groups: Ptf1a-only, Sox2-only and Atoh1-only both at E12 (I-M) and E13 (N-R). ***p < 0.001, ****p < 0.0001.

### Notch1 regulates inhibitory versus excitatory precursor fate decision in Sox2^+^ ECPs

First, we asked whether the choice between Ptf1a^+^ and Atoh1^+^ cells requires Notch signaling. We examined Notch1 loss of function (LOF) either in clones using *Sox2^CreERT2^/Notch1^flox^/Atoh1^GFP^* conditional knockout (KO) mice (Figure S6A-S6E), or in Presenilin1 KO (Psn1) mice, which have a near-complete loss of Notch signaling activity (Figure S6F-S6H). Because of the mutual suppression of Ptf1a and Atoh1, the wild type cerebellar precursors rarely co-express Atoh1 and Ptf1a (Yamada et al., 2014). However, bi-potential Sox2^+^ ECPs may briefly express both fate potentials before a decision is made. If so, we might find a snap shot of this moment in their development in the scRNAseq data. Examining these data at E10-E13, we did indeed find Sox2^+^/Atoh1^+^/Ptf1a^+^ triple positive cells, demonstrating the existence of individual bi-potential ECPs at the gene regulatory level. We found that these cells are very rare between E10 and E11, quickly peaking between E11 and E12 before dropping again between E12 and E13 (Figure 5A). The peak at E12 indicates that this is a critical time point for the fate choice of Sox2^+^ ECPs, we therefore examined Ptf1a and Atoh1 protein expression in Notch LOF conditions at E12. We found that in control ECP clones (Figure 5B-5D” and 5H) and control PS1^+/-^ mice (Figure 5I-5K” and 5O), the vast majority of committed precursors exclusively express either Ptf1a or Atoh1, with very few cells expressing detectable levels of both markers. In contrast, in Notch1^-/-^ clones (Figure 5E-5G” and 5H) and in PS1 KO mice (Figure 5L-5N” and 5O) we observed a significant ∼3.5-fold increase in the number of double-positive precursors. Interestingly, most of these cells were found at the VZ/RL boundary extending into the VZ. Together with the lineage tracing evidence, these data suggest that individual daughters of ECPs can adopt either an inhibitory or an excitatory fate and that the decision between these two fates requires Notch signaling. If this is correct, perturbing Notch activity should perturb the ratio of inhibitory to excitatory precursors.

**Figure 5.**
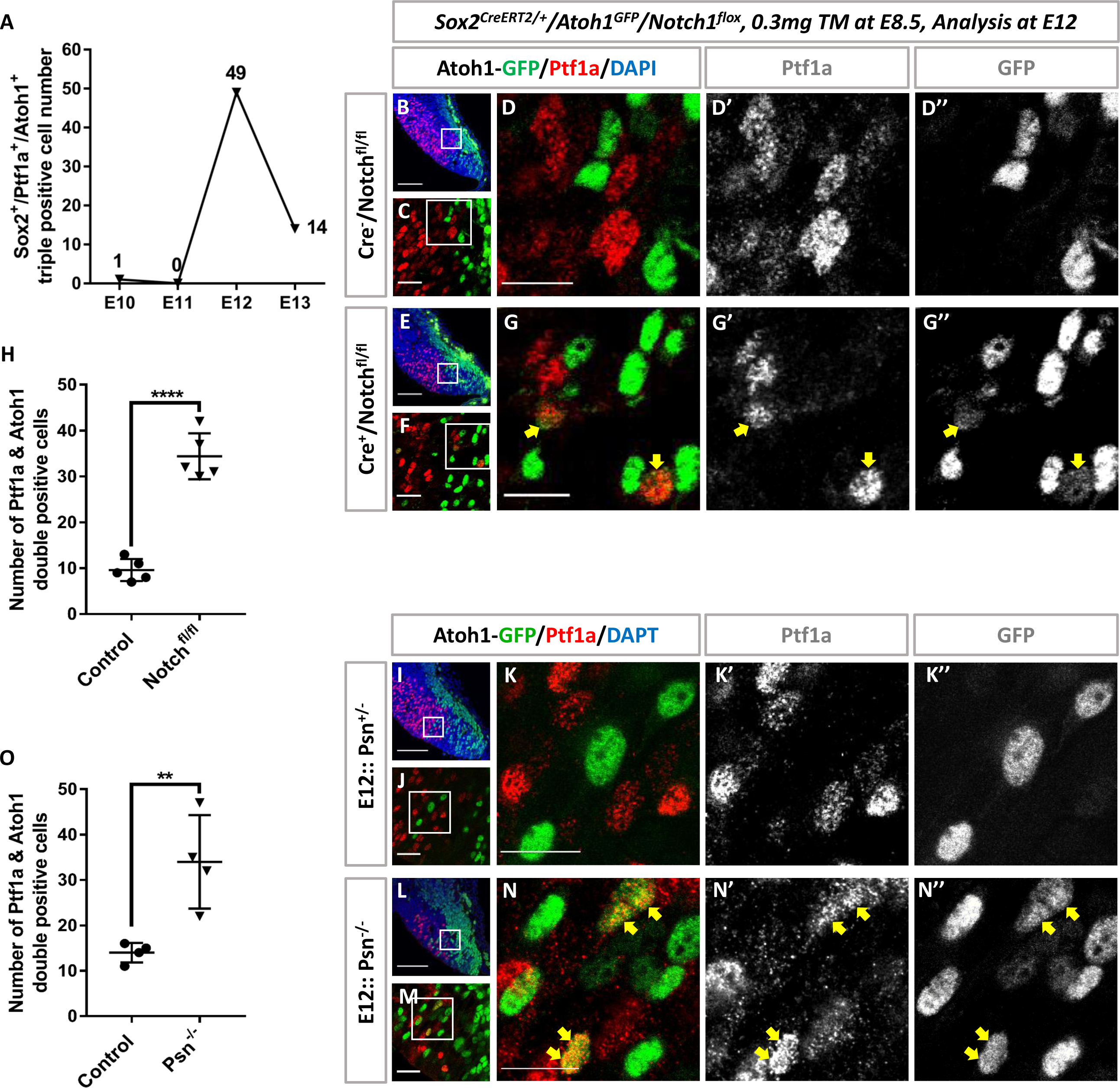
Notch loss of function increases Ptf1a and Atoh1 double positive cells at the VZ-RL boundary: (A) Analysis of Sox2^+^/Ptf1a^+^/Atoh1^+^ triple positive cells in scRNAseq data from embryonic cerebellar progenitors. (B and C) Immunostaining for Ptf1a^+^ (red) and Atoh1-GFP^+^ (green) reveals very few double positive cells at E12 in control cerebella. n=5. (D-D’’) Higher magnification of the rectangular region in (C). (E and F) Immunostaining for Ptf1a^+^ (red) and Atoh1-GFP^+^ (green) reveals increased double positive cells at E12 in Notch1 cKO cerebellum. n=5. (G-G’’) Higher magnification of the rectangular region in (F). Arrows indicate Ptf1a^+^ and Atoh1-GFP^+^ double positive cells. (H) Percentage of Ptf1a^+^ and Atoh1-GFP^+^ double positive cells in Control and Notch1 cKO cerebella. (I and J) Immunostaining for Ptf1a^+^ (red) and Atoh1-GFP^+^ (green) double positive cells at E12 in Control cerebellum. n=4. (K-K’’) Higher magnification of the rectangular region in (J). (L and M) Immunostaining for Ptf1a^+^ (red) and Atoh1-GFP^+^ (green) double positive cells at E12 in Psn KO cerebellum. n=4. (N-N’’) Higher magnification of the rectangular region in (M). (O) Percentage of Ptf1a^+^ and Atoh1-GFP^+^ double positive cells in Control and Psn KO cerebella. Arrows indicate Ptf1a^+^ and Atoh1-GFP^+^ double positive cells. Nuclei were stained with DAPI (blue). Scalebars=100 µm, 25 µm and 15 µm. Data presented as mean ± SD. **p < 0.01, ****p < 0.0001.

To test this, we quantified the numbers and ratios of Atoh1^+^ and Ptf1a^+^ cells under Notch1 loss of function conditions. First, we quantified the number of Sox2^+^ cells that also express Atoh1 and found a progressive increase in the number of these cells with the progressive reduction in number of Notch1 copies (Figure 6A-6D). Furthermore, ectopic Sox2^+^/Atoh1^+^ cells were found within the VZ especially under Notch1^-/-^ conditions (Figure 6C), indicating a possible conversion of inhibitory VZ precursors into excitatory precursors. To directly test whether this is the case, we quantified the relative abundance of Ptf1a^+^ and Atoh1^+^ cells and found that the increase in Atoh1^+^ cells came at the expense of Ptf1a^+^ cells which were significantly decreased in Notch1^-/-^ ECP clones (Figure 6G-6I) as well as Psn1 KO mice (Figure 6L-6N). We observed a similar change in the relative expression of Atoh1 and Ptf1a mRNAs (Figure 6E-6F, 6J-6K, S7A and S7B). Importantly, the ratio of the increase in Atoh1^+^ cells was nearly identical to the ratio of decrease in Ptf1a^+^ cells (0.672 vs 0.673, Figure 6I and 6N), further supporting a Notch-dependent common origin of the two precursors. To confirm that this is a reduction in inhibitory GABAergic precursors and not only in Ptf1a expression, we examined the expression of other VZ precursor markers, namely Olig2, which maintains the identity of PC progenitors (Ju et al., 2016), and Lhx1/5, a marker for early GABAergic cells (Hori et al., 2008; Muguruma et al., 2015), in Psn1 KO mice and found that levels of Olig2 mRNA were also decreased (Figure S7C) as were the numbers of Olig2^+^ and Lhx1/5^+^ cells, concomitantly with the increase in the number of Atoh1^+^ cells (Figure S7D-S7H). Importantly, these changes in cell fate were not accompanied by changes in the degree of proliferation nor cell death (Figure S8). Therefore, Notch1 signaling regulates the choice between excitatory and inhibitory precursor fate. Finally, we asked whether Notch inhibition also suppresses GABAergic fate in human cerebellar organoids by treating differentiating organoids with the gamma-secretase inhibitor Dibenzapine (DBZ) which results in a strong inhibition of Notch signaling. This resulted in a strong increase of Atoh1 expression and reduction of both Ptf1a expression and Calbindin^+^ cell number, indicating a loss of GABAergic PCs (Figure S9).

**Figure 6.**
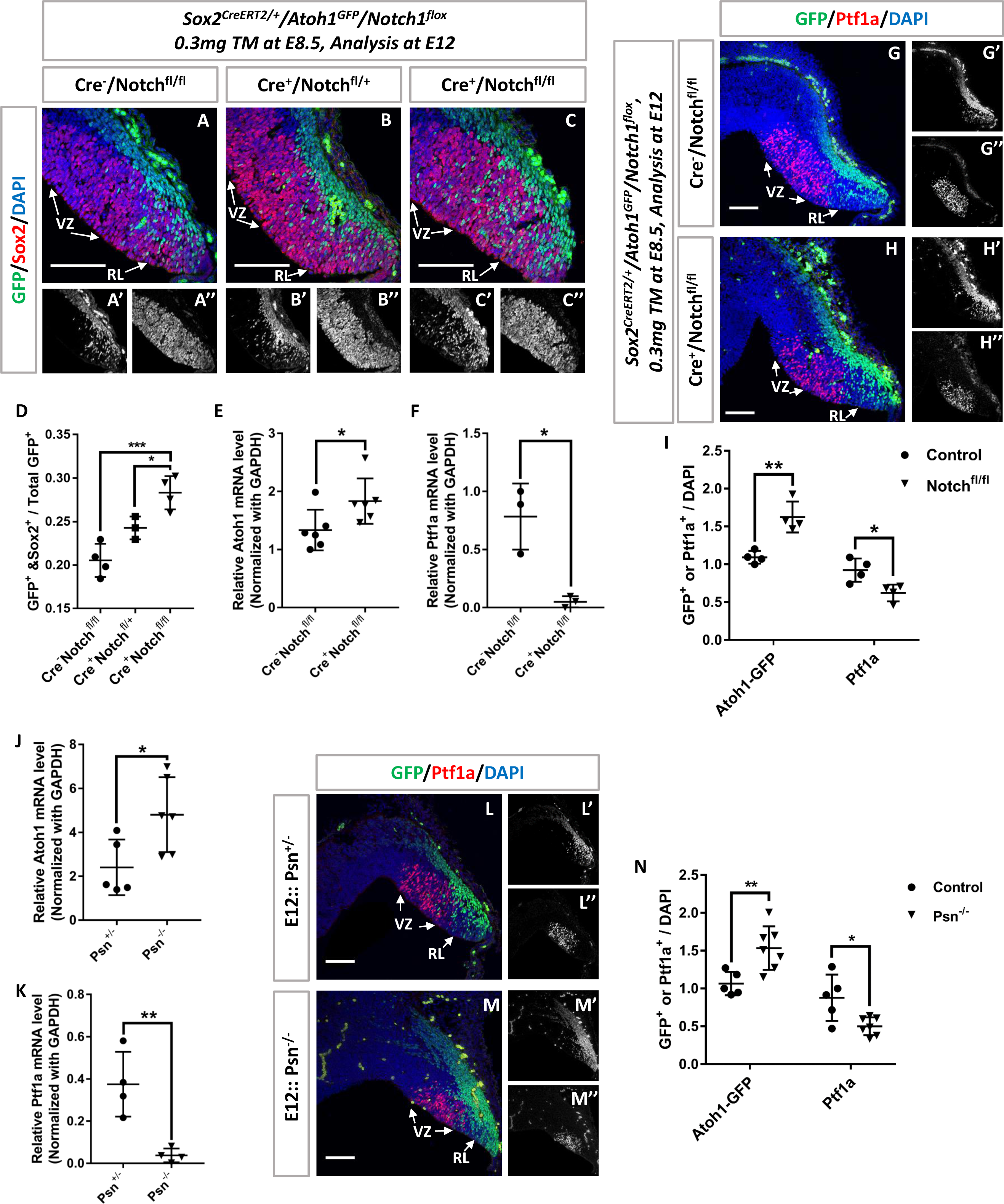
Notch loss of function in cerebellar progenitors favors Atoh1^+^ progenitors at the expense of Ptf1a^+^ progenitors: (A-C’’) Immunostaining for Sox2^+^ (red) and Atoh1-GFP^+^ (green) at E12 in Cre^-^ /Notch^fl/fl^ (n=4), Cre^+^/Notch^fl/+^ (n=3) and Cre^+^/Notch^fl/fl^ (n=4) mice. (D) Percentage of Atoh1-GFP^+^/Sox2^+^ double-positive cells within all Atoh1-GFP^+^ cells in the three genotypes. (E and F) Comparison of Atoh1 (n=6) and Ptf1a (n=3) mRNA levels in Cre^-^/Notch^fl/fl^ and Cre^+^/Notch^fl/fl^ mice using RT-PCR at E12. (G-H’’) Immunostaining for Ptf1a^+^ (red) and Atoh1-GFP^+^ (green) at E12 in control cerebella (G-G’’) and in Notch1 conditional KO (cKO) cerebella (H-H’’). n=4.(I) Percentage of Ptf1a^+^ cells or Atoh1-GFP^+^ cells among all cells (DAPI, blue) in control and Notch1 cKO cerebellum. (J and K) Comparison of Atoh1 (n=5 for Control group and n=6 for Psn KO group) and Ptf1a (n=4) mRNA levels in Control and Psn KO cerebellum using RT-PCR at E12. (L-M’’) Immunostaining for Ptf1a^+^ (red) and Atoh1-GFP^+^ cells (green) at E12 in Control (L-L’’, n=5) and Presenilin1 (Psn) KO cerebella (M-M’’, n=7). (N) Percentage of Ptf1a^+^ cells or Atoh1-GFP^+^ cells within all cells (DAPI) in control and Psn KO cerebella. Scalebars=100 µm and 25 µm. Data presented as mean ± SD. *p < 0.05; **p < 0.01; ***p < 0.001.

### Notch gain of function in ECPs maintains their stem cell fate in cerebellar primordium

During cortical development, transient gain of Notch function results in progenitors skipping the production of early born (deep layer) neurons and thus favoring the production of late-born (upper layer) cells. Consequently, the effect of Notch gain of function is to maintain cells in progenitor state to contribute to the temporal axis of neuronal diversification. However, our data show that Sox2^+^ ECPs generate inhibitory and excitatory lineages in the cerebellum simultaneously and that the inhibitory versus excitatory binary fate choice requires Notch signaling. Gene expression analysis suggests that ECPs have the highest levels of Notch activity followed by inhibitory VZ precursors with excitatory RL precursors having very low Notch signaling activity (Figure 4). To better understand the role of Notch signaling in the balance between maintaining progenitors and regulating binary cell fate, we analyzed the effects of overexpression of the constitutively active Notch1 intracellular domain (NICD) on ECPs. We generated Notch1 gain of function (GOF) ECP clones using *Sox2^CreERT2/+^/ Gt(ROSA)26Sor^tdTomato^/R26R^stop-NICD-nGFP^* mice, and examined cell fate inside the clones at E12, when all three cell types (ECPs, VZPs and RLPs) are abundant. We found that Notch GOF strongly reduces the number of both Ptf1a^+^ and Atoh1^+^ precursors (Figure 7A-7K, S10A-S10D’’ and S10G). However, whereas we were able to detect Ptf1a^+^ cells (Figure 7C-7D” and S10A-S10B’’), no Atoh1^+^ cells were detected within the Notch1 GOF clones (Figure 7H-7I” and S10C-S10D’’). Quantification confirmed that the effect was significantly stronger on Atoh1^+^ cells than on Ptf1a^+^ cells (Figure 7K and S10G). Lastly, we checked Sox2 expression within the NICD-GFP clones and found that Notch GOF in progenitor cells significantly increased Sox2^+^ cells both in the RL and VZ (Figure 7L-7P, S10E-S10F’’ and S10H) at the expense of Ptf1a^+^ and Atoh1^+^ cells. Importantly, the inhibition of differentiation induced by Notch GOF was long lasting (Figure S11). Almost all NICD-GFP^+^ cells retained an undifferentiated state at E16 with very few NICD-GFP^+^ cells differentiating into Calbindin^+^ PCs (Figure S11A-S11C’’), and none of the cells differentiating into Atoh1^+^ GC precursors (Figure S11D-S11F’’), while both in the RL and VZ, the NICD-GFP^+^ cells retained high levels of Sox2 expression (Figure S11G-S11L). Altogether, our data demonstrate that Notch activity dictates cell fate in bi-potential cerebellar progenitor cells.

**Figure 7.**
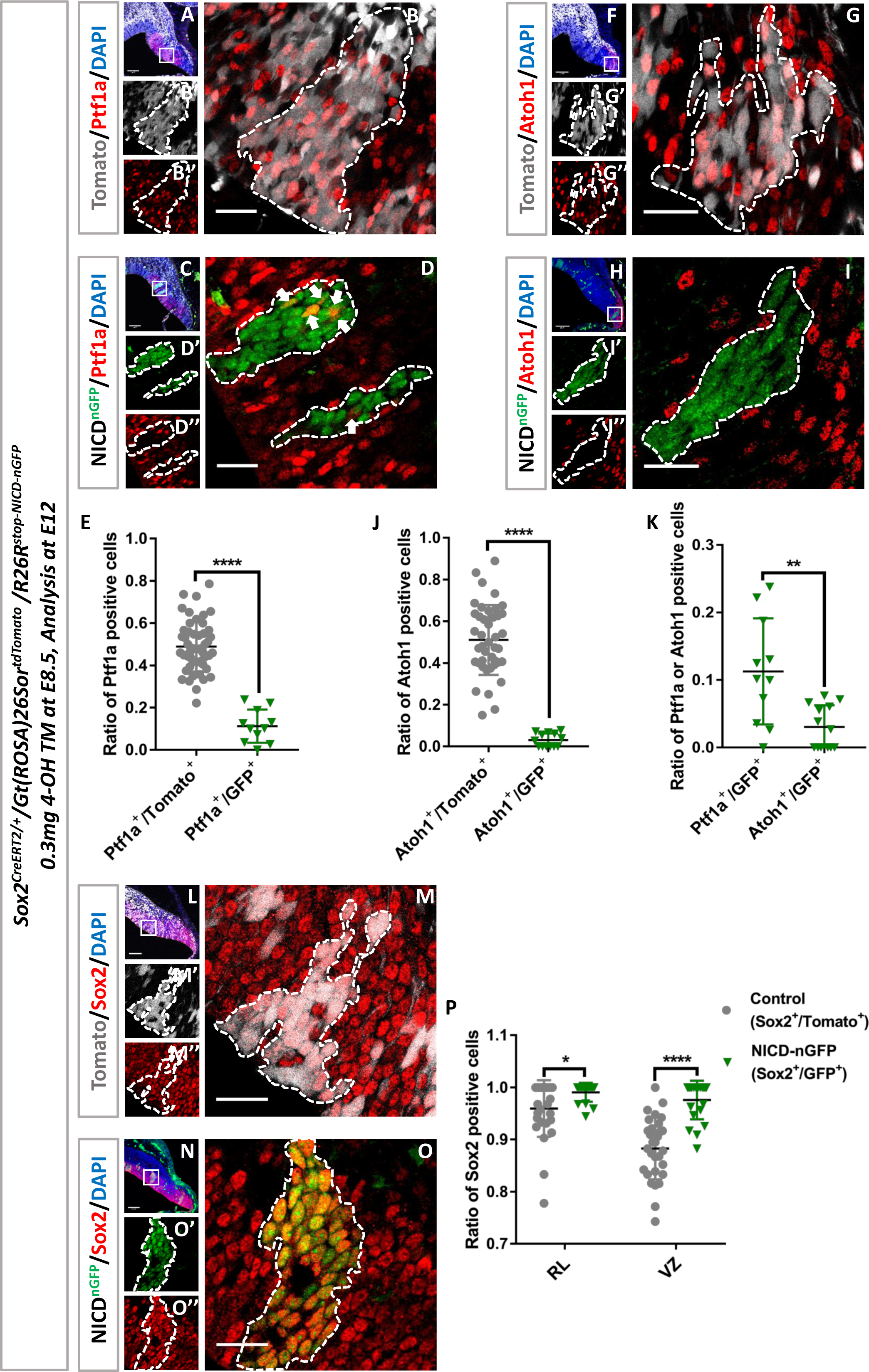
Notch GOF in ECPs inhibits their differentiation: (A) Immunostaining for Ptf1a^+^ (red) in tdTomato^+^ wild-type clones (grey) at E12. n=4 embryos, totally 51 clones in the VZ. (B-B’’) Higher magnification of the rectangular region in (A). (C) Immunostaining for Ptf1a^+^ (red) in NICD-GFP^+^ clones (green) at E12. n=4 embryos, totally 11 clones in the VZ. (D-D’’) Higher magnification of the rectangular region in (C). Arrows indicate NICD-GFP^+^/Ptf1a^+^ double-positive cells. (E) Percentage of Ptf1a^+^ cells in control (tdTomato^+^) versus NICD-GFP^+^ VZ clones. (F) Immunostaining for Atoh1^+^ (red) in tdTomato^+^ wild-type clones (grey) at E12. n=4 embryos, totally 42 clones in the RL. (G-G’’) Higher magnification of the rectangular region in (F). (H) Immunostaining for Atoh1^+^ (red) in NICD-GFP^+^ clones (green) at E12. n=7 embryos, totally 13 clones in the RL. (I-I’’) Higher magnification of the rectangular region in (H). (J) Percentage of Atoh1^+^ cells in control (tdTomato^+^) versus NICD-GFP^+^ RL clones. (K) Comparison of the ratios of Ptf1a^+^ cells and Atoh1^+^ cells in NICD-GFP^+^ clones. (L) Immunostaining for Sox2^+^ (red) in tdTomato^+^ wild-type clones (grey) at E12. n=4 embryos, totally 28 clones in the RL and 30 clones in the VZ. (M-M’’) Higher magnification of the rectangular region in (L). (N) Immunostaining for Sox2^+^ (red) in NICD-GFP^+^ clones at E12. n=4 embryos, totally 15 clones in the RL and 22 clones in the VZ. (O-O’’) Higher magnification of the rectangular region in (N). (P) Percentage of Sox2^+^ cells in control (tdTomato^+^) versus NICD-GFP^+^ RL and VZ clones. Nuclei marked with DAPI (blue). Scalebars=100 µm and 25 µm. Data presented as mean ± SD. *p < 0.05; **p < 0.01; ****p < 0.0001.

## DISCUSSION

In this work we show that, in contrast to what is classically described, GABAergic and glutamatergic neurons of the cerebellar cortex in both mouse and human organoids can indeed derive from common progenitors which we here term Embryonic Cerebellar Progenitors (ECPs). Population and single cell lineage tracing approaches demonstrated that Sox2^+^ ECPs cells are likely to give rise to multiple cerebellar cell types, including GABAergic and glutamatergic neurons as well as astroglial cells. These progenitors span the RL and the VZ and are characterized by the expression of Sox2^+^ and very high levels of Notch activity. This finding led us to ask how these common progenitors give rise to different types of neurons. Analysis of scRNAseq data suggested cell fate diversification may be influenced by Notch signaling. This was surprising because while Notch activity is known to distinguish progenitors from differentiated daughters in all lineages where it was examined, no evidence for a role of Notch signaling in neuronal binary cells fate choice has been found in mammalian brain neurogenesis. Our *in vivo* genetic analyses using loss and gain of Notch1 function, as well as Psn1 KO mice, support a simple model whereby early common progenitors generate precursors that express different levels of the Notch ligands Delta1 and Delta3 and signal back to their mother cells and to each other to segregate into different cell fates. This creates a binary choice where different types of precursors eventually generate excitatory versus inhibitory cells, respectively. The expression of Delta3 largely by RL precursors allows these precursors to further repress Notch activity cell autonomously, as Delta3 is thought to act as a cell autonomous cis-inhibitor of the Notch receptor (Ladi et al., 2005). We thus provide mechanistic evidence for how common progenitors can generate both excitatory and inhibitory lineages in the mammalian brain. It will be interesting to test precisely how Notch activity in ECPs or their daughters plays a role in disorders of cell fate or stem cell behavior such as medulloblastomas, as recent work suggests (Ballabio et al., 2020b).

It is very interesting to note that while PCs and GCs themselves are generated on very different temporal scales, with PCs born much earlier than GCs, their precursors are specified at the same time. GC production is late because GC precursors undergo several rounds of transient amplification before generating neurons, thus creating this pseudo-heterochrony. The transient amplification of GC precursors is driven by Atoh1 which itself regulates, and is regulated by, Notch signaling. Our work provides a developmental and molecular framework for how common progenitors can create cell type diversity across different time scales through the highly-conserved process of binary cell fate specification. In *Drosophila* neuronal lineages, the two daughter neurons of the same precursor use Notch to adopt different identities during terminal cell division (Artavanis-Tsakonas et al., 1999; Bhat, 2014; Pinto-Teixeira and Desplan, 2014). Similarly, spinal cord inhibitory interneurons use Notch activity to adopt different identities (Peng et al., 2007). Single cell analysis in the neocortex is beginning to reveal significant diversity of excitatory neuronal fates in the same cortical layers (Pfeffer and Beltramo, 2017). It would be interesting to test whether Notch signaling also regulates the fate of cortical sister neurons where the lack of sufficient markers of neuronal diversity may have precluded such findings in the past.

In *Drosophila*, part of the generation of cells with different Notch-dependent fates relies on asymmetric cell division of progenitors and the biased segregation of Notch inhibitors. Whether the production of Atoh1^+^ and Ptf1a^+^ cells is similarly modulated by asymmetric division of Sox2^+^ progenitors needs further investigation. Because the bone morphogenetic proteins (BMPs) signaling and the diffusible mitogen Sonic hedgehog (Shh) signaling are also key regulators involved in GABAergic and glutamatergic lineage generation and differentiation (Fleming et al., 2013; Huang et al., 2010; Laudet et al., 1993; Machold et al., 2007), the integration between these activities and binary cell fate determination would be an interesting avenue of future work. Recent work suggests a common developmental basis for the evolution of cerebellar nuclei from conserved cell types (Kebschull et al., 2020). The derivation of at least some GCs and PCs from common progenitors via Notch-mediated binary cell fate choice suggests one possible genetic basis for how the excitatory to inhibitory ratio may change during evolution.

## STAR✰METHODS

Detailed methods are provided in the online version of this paper and include the following:

• KEY RESOURCES TABLE

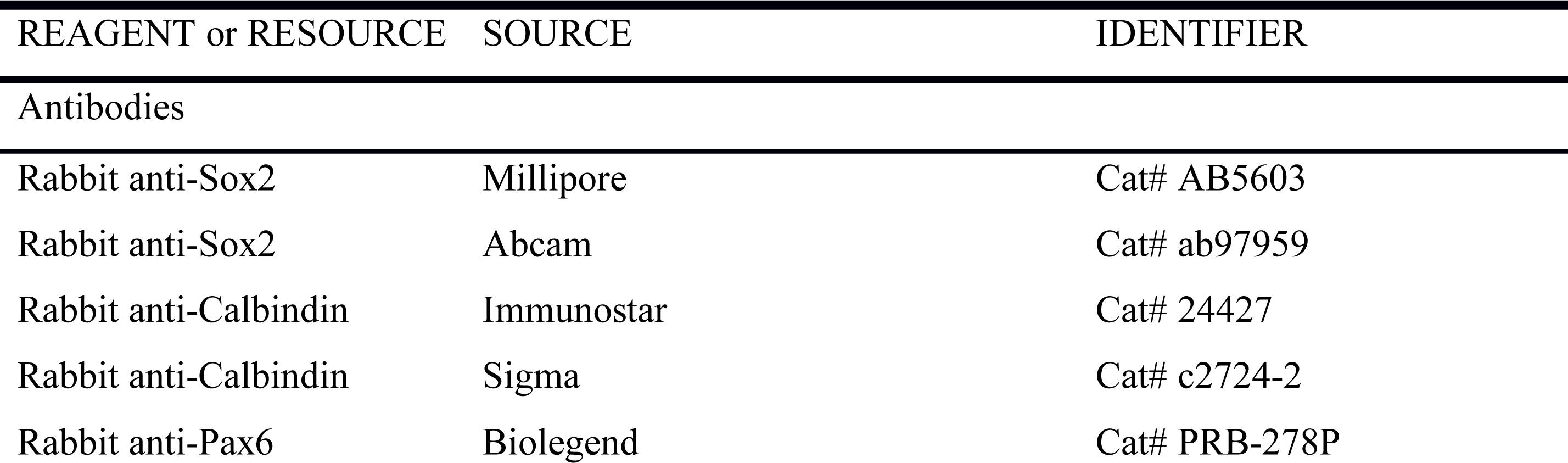

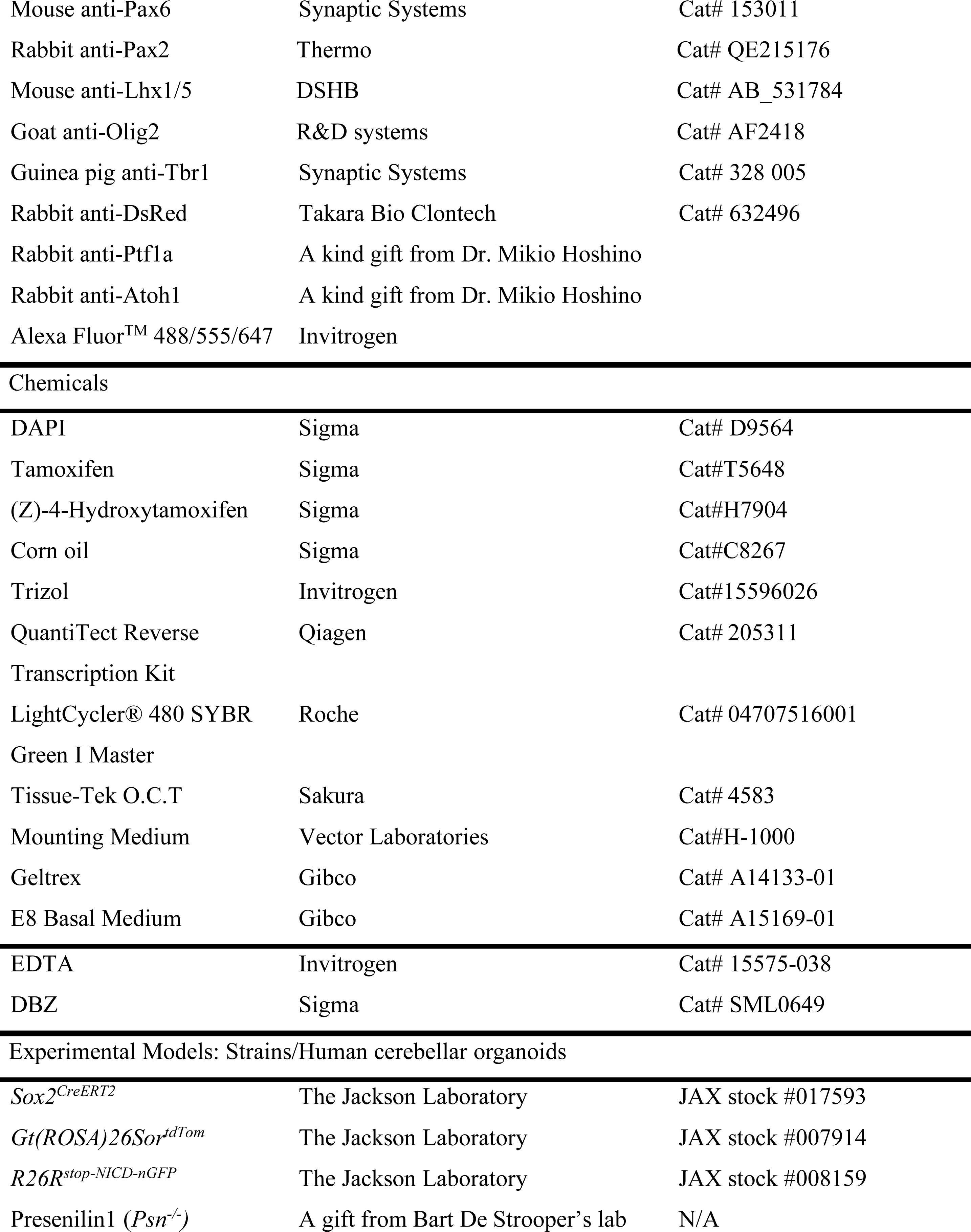

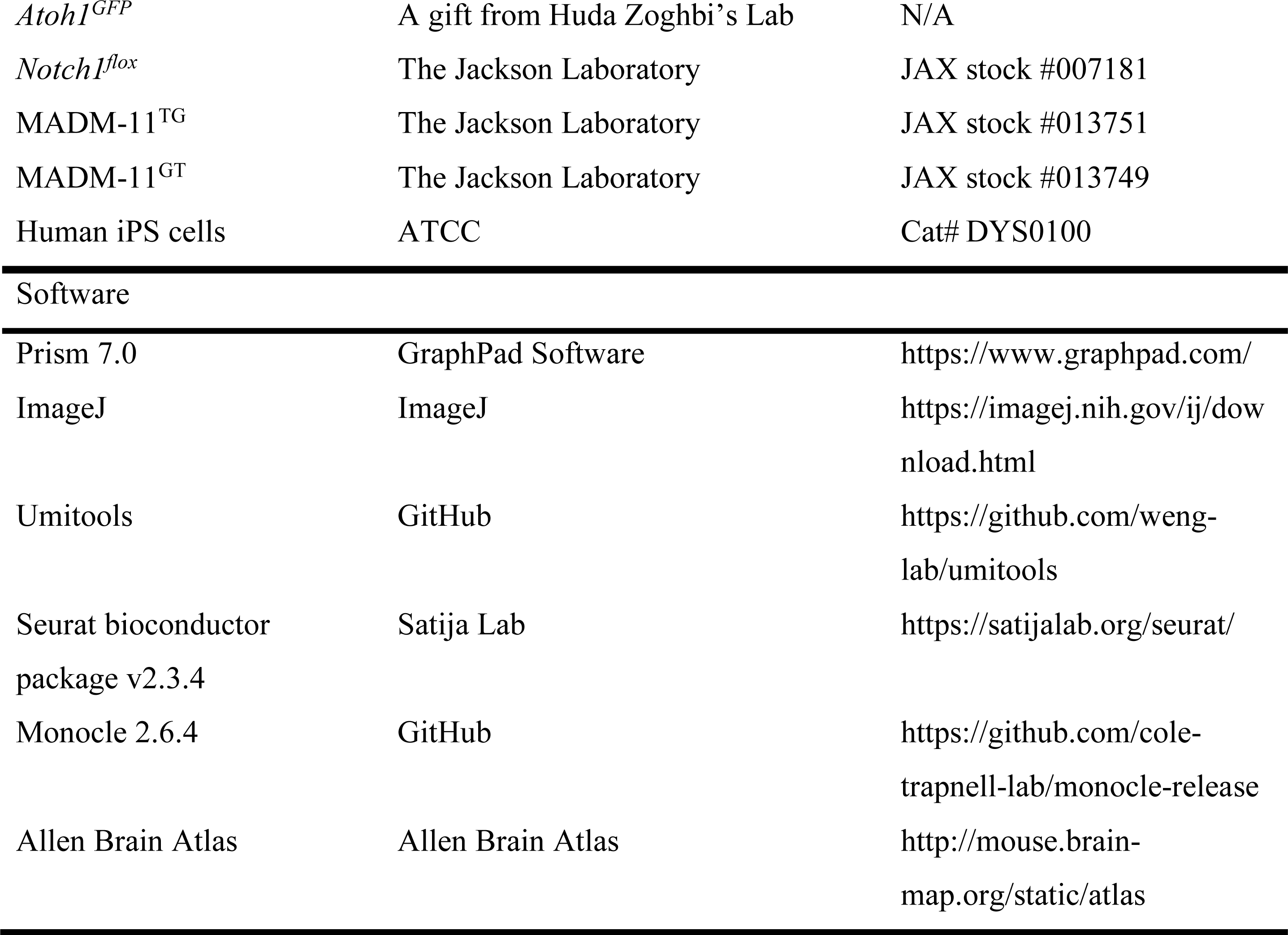

### Mice

All animal experiments in this study were carried out in accordance with animal welfare regulations and have been approved by Ethic Committee and French regulatory authorities of the respective institutes. The *Sox2^CreERT2^* mice were crossed with *Gt(ROSA)26Sor^tdTom^* reporter mice, and then crossed with *Atoh1^GFP^* mice to generate the lineage tracing line. Notch gain-of-function mice were generated by breeding *Sox2^CreERT2^/ Gt(ROSA)26Sor^tdTom^* with *R26R^stop-NICD-Ngfp^* (Murtaugh et al., 2003). Presenilin1 deficient (*Psn^-/-^*) mice crossed with *Atoh1^GFP^* mice to get *Atoh1^GFP^/Psn^-/-^* mice. And Notch conditional knock out mice was generated by crossing *Notch1^flox^* with *Sox2^CreERT2^/ Atoh1^GFP^* mice. Mice were genotyped genomic DNA isolated from toe tissue. The primers used for genotyping were shown in table S2.

### Tamoxifen Administration

Tamoxifen (TM, Sigma) or 4-Hydroxytamoxifen (4-OH, Sigma) was dissolved to a final concentration of 1mg/ml or 3mg/ml in 90% corn oil (Sigma) with 10% ethonal (Sigma). For *Sox2^CreERT2^/ Gt(ROSA)26Sor^tdTom^ / Atoh1^GFP^* mice, if collected the samples at embryonic stages, 0.1mg TM was intraperitoneal (i.p.) injected into per pregnant female at E9.5 or E10.5. If collected the samples at postnatal stages, each pregnant female was injected 0.03mg TM (i.p.) at E10.5. For *Sox2^CreERT2^/Gt(ROSA)26Sor^tdTom^/R26R^stop-NICD-nGFP^* mice and *Sox2^CreERT2^/ Notch1^flox^* mice, 0.1ml 4-OH TM or TM (3mg/ml) was injected into the pregnant females (i.p.) at E8.5, respectively.

### RNA extraction and Real time PCR (RT-PCR)

Total RNA was isolated from whole cerebellar tissue samples at E13 using Trizol Reagent (Invitrogen). The RNA concentration was measured by a spectrophotometer (NanDrop1000; Thermo) followed by a reverse transcription process using PrimeScript RT reagent kit (Roche). Real time PCR analysis was performed using SYBR green mix (Roche) and the values were normalized to β-actin values. The primer pairs for different genes were shown in table S3. PCR conditions used here were denaturing at 95 °C for 10 s, followed by 40 cycles of 95 °C for 5 s and 60 °C for 30 s. Data were analyzed using the comparative threshold cycle (Ct) method, and results were expressed as fold difference normalized to GAPDH.

### Immunohistochemistry and antibodies

For the samples collection, embryos before E13.5 were fixed in 2% paraformaldehyde (PFA) in PBS at 4 °C for 2-3 hours, and embryos after E13.5 but before born first were perfused with 2% PFA, then post-fixed for 24 hours. Whereas, samples were collected at postnatal stages, perfused the mice with 1 X PBS, followed by 4% PFA perfusion, and then post-fixed in 4% PFA for another 24 hours. Dehydrated embryos or the whole head in 30% sucrose in 1 X PBS overnight (o/n). After all the samples sank into the bottom of the tube, embedded them in OCT compound (TissueTek) and frozen at −20 °C. Sagittal sections were made by cryostat (Leica) at 20 µm and then stored slices at −80 °C. For the immunostaining, sections were fixed with 4% PFA for 10 minutes at room temperature (RT), then blocked with 10% normal donkey or goat serum in 1 X PBS with 0.1% Triton (PBT) for 1 hour at RT followed by 3 times washing in 1 XPBT. Thereafter, these sections were incubated with primary antibodies diluted in 0.1% 1 X PBT containing 1% normal donkey or goat serum o/n at 4 °C or 3-4 hours at RT. After 3 times washing with 1 X PBT, incubated with appropriate secondary antibodies conjugated with Alexa Fluor 488, Alexa Fluor 555, or Alexa Fluor 647 (1:500, Invitrogen) in 0.1% 1 X PBT containing 1% normal donkey or goat serum for 1-2 hours at RT. Washed with 1 X PBT for 3 times, then counterstained the slides with DAPI (1:2000, Sigma) and mounted by using Vectashield (Vector) after rinsing. Antigen retrieval was performed by using 10 mM sodium citrate buffer, pH 6.0, boiled 5 minutes in microwave, and cooled down in RT for about 20 minutes for Lhx1/5 and Olig2 staining. Primary antibodies used in this study were rabbit anti-Sox2 (1:500, Millipore, AB5603), rabbit anti-Ptf1a and Rabbit anti-Atoh1 (1:200, a kind gift from Dr. Mikio Hoshino, National Center of Neurology and Psychiatry · Department of Biochemistry and Cellular Biology, Japan), rabbit anti-Calbindin (1:500, Immunostar, 24427), rabbit anti-Pax6 (1:300, Biolegend: PRB-278P), mouse anti-Pax6 (1:300, Synaptic Systems: 153011), rabbit anti-Pax2 (1:200, Thermo, QE215176), mouse anti-Lhx1/5 (1:100, DSHB, AB_531784), goat anti-Olig2 (1:500, R&D systems, AF2418) and Guinea pig anti-Tbr1 (1:500, Synaptic Systems, 328 005). After staining, images were obtained by using confocal microscope (Olympus FV-1200 or Leica SP8).

### Primary culture of mouse cerebellar progenitor cells

#### Preparation

Two days before cell isolation, 13mm cover slips were placed in a 24 well plate, then the plate was coated with 500 ul poly-L-ornithine hydrobromide (Sigma, P4957) and incubated overnight. Next, the plate was washed 3 times with sterile water, then coated with 500 ul laminin (Final conc: 5ug/ml, Sigma, L2020) and incubated overnight.

#### Isolation and culture of cerebellar progenitor cells

Cerebellar progenitors were harvested from *Sox2^CreERT2^/ Gt(ROSA)26Sor^tdTomtato^* pregnant mice. In brief, cerebella were dissected from E11.5 mouse embryos in L15 medium (Gibco, 11415064) and enzymatically dissociated to single cells using 0.05% trypsin/EDTA (Gibco, 25300-054) plus 30% SVF (Invitrogen, 10270106) plus DNAse (Serlabo, LS002138). The cells were collected and resuspended in proliferation medium: Neurobasal medium without phenol red (Gibco, 12348-017) containing B27 supplement (Gibco, 17504-044), L-glutamax (Gibco, 35050-061), 20 ng/ml mouse epidermal growth factor (EGF, Thermofisher, PMG8041), 10 ng/ml mouse basic fibroblast growth factor (bFGF, Thermofisher, PMG0035), 1M HEPES (Gibco, 12509079) and insulin (Sigma, I0516). Cells were plated on coverslips in a precoated 24-well plate with 0.05 ng/ml Tamoxifen treatment, then cultured at 37 °C with 5% CO_2_ in an incubator (Thermo Scientific).

#### Time-Lapse Video Recording

Primary cultured progenitor cell proliferation was tracked using time-lapse video recording. Two hours after cell seeding, the 24-well plate was transferred to a videomicroscope (Zeiss AxioObserver 7) with a humidified incubator at 37 °C with a constant 5% CO_2_ supply. Timlapse images for both Cy3 and bright field were acquired every 30 min, 45 min and 60 min for for 72 hours, day 4 and day 5 and day 6-day12, respectively.

#### Induction of Purkinje and Granule cells from Sox2^+^ cerebellar progenitors

Three days after time-Lapse Video Recording, the proliferation medium was changed to differentiation medium as described previously (Kawasaki et al., 2000; Su et al., 2006). First, to induce Pax6^+^ granule neurons, cells were cultured in α-MEM medium (Gibco, 12561056) supplemented with 5% KSR (Gibco, 10828010), 0.1mM 2-ME (Gibco, 31350010) and 0.5 nM mouse BMP4 (R&D, 5020-BP-010). Three days later, the medium was changed to Neurobasal-Plus (Gibco, A3582901) supplemented with B27-Plus (Gibco, A3582801), Ascorbic acid (Sigma, A4403) and 0.5 nM mouse BMP4 in order to induce the production of Calbindin^+^ Purkinje neurons. At the end of the 12 days of recording, cells were fixed directly by using 4% PFA and immunostaining was performed.

#### Cell counts

Confocal images for single layer scanning of sagittal cerebellar sections were calculated for each developmental stage (E11.5, E12 and E13) after DAPI staining. Each section took the average of the four values that obtained from 4 single layer calculating, which could form a Z stack. And each cerebellar sample counted 6-8 sections that took from the beginning to the end of the cerebellum. All quantifications were done blinded to the genotyping. For each stage littermates were analyzed and all groups of quantifications were carried out from at least 3 individuals.

#### scRNAseq quantification and statistical analyses

Aligned 10X data were retrieved from ENA: PRJEB23051 data set for the following samples: E10, E11, E12 and E13. Umitools has been used to generate gene-cell matrices with the following parameters: --extract-umi-method=tag, --umi-tag UB, --cell-tag CB, --per-gene, --gene-tag GX, -- per-cell. Genes not expressed in any cells were removed from considerations, as were all mitochondrial and ribosomal protein genes. To remove likely dead or multiplet cells from downstream analyses, cells were discarded if they had less than 3500 UMIs, greater than 15000 UMIs, or were composed of over 10% mitochondrial UMIs. The final dataset was composed of 14637 cells and 18937 genes. Then Seurat bioconductor package v2.3.4(Butler et al., 2018) has been used to do cell-cell comparison and identify cell types. First, we performed a t-SNE (t-distributed stochastic neighbor embedding) with the first 20 principal components after application of PCA reduction. This allowed us to visualize the grouping of cells and the expression of genes of interest. Expression of cells in 3 populations has been represented with violin plot and differential expression between the 3 populations has been calculated with a Welch two sample t-test procedure.

Monocle 2.6.4(Qiu et al., 2017) was used to infer the pseudotime trajectory. As we worked with UMI count data, we assumed that the data were distribued among a negative binomial distribution with fixed variance. The genes that “define progress” were selected using the unserpervised procedure “dpFeature”: we first selected genes expressed in at least 5% of all the cells. We then run reduceDimension with tSNE as the reduction method, num_dim=10”, norm_method=“log” and max_components = 2. Finally, cells were clustered with the density peak clustering algorithm by setting P to 2 and Δ to 5 (and skip_rho_sigma = T to facilitate the computation). The top 1000 significantly differentially expressed genes between clusters were selected as the ordering genes. The state 3 where Sox2 is expressed and Atoh1 not expressed was defined as the start of the pseudotime. The seurat FindMarkers function was used to identify the top 10 genetic markers of each lineage state’s.

#### MADM Mouse Lines and Maintenance

MADM employs Cre recombinase/loxP-dependent interchromosomal recombination highlighting two scenarios: (i) Recombination occurs in G2 phase of the cell cycle with X segregation (G2-X MADM clone) can create two distinctly labelled daughter cell lineages from their common mother progenitor cell; (ii) Recombination occurs in G1 phase or in G2 phase followed by Z segregation (G1/G2-Z MADM clone), one or both daughter cell lineages will be labelled in yellow. Mouse protocols were reviewed by institutional ethics committee and preclinical core facility (PCF) at IST Austria and all breeding and experimentation was performed under a license approved by the Austrian Federal Ministry of Science and Research in accordance with the Austrian and EU animal laws (license numbers: BMWF-66.018/0007-II/3b/2012 and BMWFW-66.018/0006-WF/V/3b/2017).. Mice were maintained and housed in animal facilities with a 12-hour day/night cycle and adequate food/water conditions according to IST Austria institutional regulations. Mouse lines with Chr. 11 MADM cassettes (MADM-11^TG^ and MADM-11^GT^), and *Sox2-CreER* have been described previously (Arnold et al., 2011; Hippenmeyer et al., 2010). All MADM-based analyses were carried out in a mixed C57BL/6J, CD1 genetic background.

#### Generation of MADM Clones in Cerebellum and Tissue Collection

To induce MADM labeling, *MADM-11^GT/GT^/Sox2^CreER^* were crossed with *MADM-11^TG/TG^* in order to generate experimental mice *MADM-11^GT/TG^/Sox2^CreER^*. The day of observed vaginal plug was defined as E0 to monitor gestation days. Pregnant mice were injected i.p. with TM (2-3 mg/pregnant female) (Sigma) dissolved in corn oil (Sigma) at E10 or E11 to induce MADM clones. Live embryos were recovered at E18–E19 through cesarean section, fostered, and raised until further analysis. At P21 experimental MADM mice were deeply anesthetized through injection of a ketamine/xylazine/acepromazine solution (65 mg, 13 mg and 2 mg/kg body weight, respectively), and confirmed to be unresponsive through pinching the paw. Mice were perfused transcardially with 4% PFA in phosphate-buffered saline (PBS, pH 7.4). Brains were removed and postfixed o/n at 4°C to ensure complete fixation. Brains were washed with PBS, and cryopreserved with 30% sucrose solution in PBS for approximately 48 hr. Brains were then embedded in Tissue-Tek O.C.T. (Sakura). For the analysis of MADM labeling, 35µm sagittal sections were directly and consecutively collected throughout the entire cerebellum and fixed to superfrost glass slides (Thermo Fisher Scientific). Sections were washed 3 times for 5 min with PBS, followed by staining with the nuclear stain DAPI (Invitrogen). After this step the slides were again washed 3 times with PBS and embedded in mounting medium containing 1,4-diazabicyclooctane (DABCO; Roth) and Mowiol 4-88 (Roth).

#### Imaging and analysis of MADM-labeled brains

Sections were imaged using an inverted LSM800 confocal microscope (Zeiss) and processed using Zeiss Zen Blue software. Confocal images were analyzed in Photoshop software (Adobe) by manually counting MADM-labeled cells. Cerebellar areas were identified by using the Allen Brain Atlas (http://mouse.brain-map.org/static/atlas).

#### Cerebellar Organoids labeling and staining

Cerebellar organoids were culture as described by Muguruma et al and Ishida et al. (Ishida et al., 2016; Muguruma et al., 2015). Human iPS cells (iPSCs, ATCC-DYS0100) were maintained in self-renewal on a layer of geltrex (Gibco, A14133-01), in E8 Basal Medium (Gibco, A15169-01) supplemented with E8 Supplement (50X). iPSC were dissociated with EDTA (Invitrogen) 0.5mM, pH 8.0, for 3 minutes incubation, to maintain cell clusters. 6000 cell/well were seed in 96 well (Sumitomo Bakelite) in differentiation medium (see Muguruma et al., 2015). One-third dilution and a full-volume replacement with medium were performed on day 7 and 14, respectively. On day 2, the aggregates were transferred from the 96 to 6 well non-tissue treated, to cultured in Neurobasal medium with Glutamax and N2 supplement. In order to sparsely label potential Sox2^+^ human ECPs, organoids were electroporated at 25d with 20ug pCAG PiggyBac (PBase), 80 ug pPB CAG LSL Venus and 20ug pGL3-SOX2Cre (for DBZ experiment); with 80 ug pPB CAG LSL Venus and 20ug pGL3-SOX2Cre (for cell 24h experiment). Organoids were transferred inside the Electroporation cuvettes (VWR, ECN 732-1136, 2mm) resuspended in Buffer 5 (under patent) and electroporation was performed with the Gene Pulser XcellTM.

The plasmid encoding an hyperactive form of the piggyBac transposase (pCMV HAhyPBase, pPBase) was a gift from https://www.sanger.ac.uk/ (Yusa et al., 2011). The piggyBac donor plasmid pPB CAG Venus were previously described (Ballabio et al., 2020a). The pSox2-cre plasmid was generated by cloning the cre coding sequence into pGL3-Sox2 (Addgene Plasmid #101761). The plasmid was used as backbone to insert before the start codon a loxP-STOP-loxP cassette by PCR, generating pPB CAG LSL Venus. DBZ experiment: organoids were treated from days 27 to 31 of differentiation, with 1 dose of 10 µM DBZ (Notch inhibitor) or DMSO, and then changed the medium at 31 d. Organoids were fixed at 26d or 41d of differentiation with 4% PFA, cryoprotected in 20% sucrose and embedded in Frozen Section Compound (Leica, 3801480). Organoids were cryosectioned at 40 µm with Leica CM 1850 UV Cryostat. Immunofluorescence staining were performed on glass slides. Blocking and antibody solutions consisted of PBS supplemented with 3% goat serum, 0.3% Triton X-100 (Sigma). Primary antibodies were incubated overnight at 4°C and secondary antibodies for 1 hour at RT. Primary antibodies used were Chicken anti-GFP (1:2000, abcam, ab13970), Rabbit anti-Sox2 (1:500, abcam, ab97959), Mouse anti-Pax6 (1:100, SantaCruz, sc-53108)and Rabbit anti-Calbindin (1:500, Sigma, c2724-2). Nuclei were stained with 1 µg/ml DAPI (Sigma). Sections and coverslips were mounted with Permanent Mounting Medium.

#### Statistical analysis

Statistical analyses were performed using GraphPad Prism software (GraphPad Software Inc., La Jolla, CA, USA). N values refers to independent animals and are detailed in the figure legends. All data are presented as mean ± SEM. Statistical testing was based on t-tests or one-way ANOVA, followed by Tukey’s honest significant test.

## ACKNOWLEDGMENTS

This work was supported by the program “Investissements d’avenir” ANR-10-IAIHU-06, ICM, an Allen Distinguished Investigator Award and the Roger De Spoelberch Foundation Prize (to B.A.H.), Armenise-Harvard Foundation, AIRC and CARITRO (to L.T.) and the European Research Council under the European Union’s Horizon 2020 research and innovation programme grant agreement No 725780 LinPro (to S.H.). T.T.Z., T.Y.L. were supported by doctoral fellowships form the China Scholarship Council and A.H.H. by a doctoral DOC fellowship of the Austrian Academy of Sciences (24812). All animal work was conducted at the PHENO-ICMice facility. The Core is supported by 2 “Investissements d’avenir” (ANR-10-IAIHU-06 and ANR-11-INBS-0011-NeurATRIS) and the “Fondation pour la Recherche Médicale”. Light microscopy work was carried out at ICM’s imaging core facility, ICM.Quant, and analysis of scRNAseq data was carried out at ICM’s bioinofmrtaics core facility, iCONICS. We thank Paulina Ejsmont, Natalia Danda and Nathalie De Geest for technical support. We are grateful to Dr. Shahragim TAJBAKHSH for providing R26R^stop-NICD-nGFP^ transgenic mice, Dr. Bart De Strooper for Presenilin1 deficient mice, Dr. Jean-Christophe Marine for *Gt(ROSA)26Sor^tdTom^* reporter mice and Dr. Martinez Barbera for *Sox2^CreERT2^ mice*. We also thanks to Dr. Mikio Hoshino for providing Atoh1 and Ptf1a antibodies. B.A.H. is an Einstein Visiting Fellow of the Berlin Institute of Health.

## AUTHOR CONTRIBUTIONS

B.A.H. and T.T.Z. designed the project and the experiments and prepared the manuscript with input from S.H. and L.T.; T.T.Z., T.Y.L. performed the majority of mouse experiments and data analysis; X.C., A.H.H., and C.S. performed the MADM experiments supervised by S.H.; N.D. helped with the preparation of MADM samples; M.A. performed the human organoid work supervised by L.T.; N.M. initiated the project. J.G. and M.B. performed bioinformatics analysis.

## DECLEARTION OF INTERESTS

The authors declare no competing interests.

---

**Figure S1.**
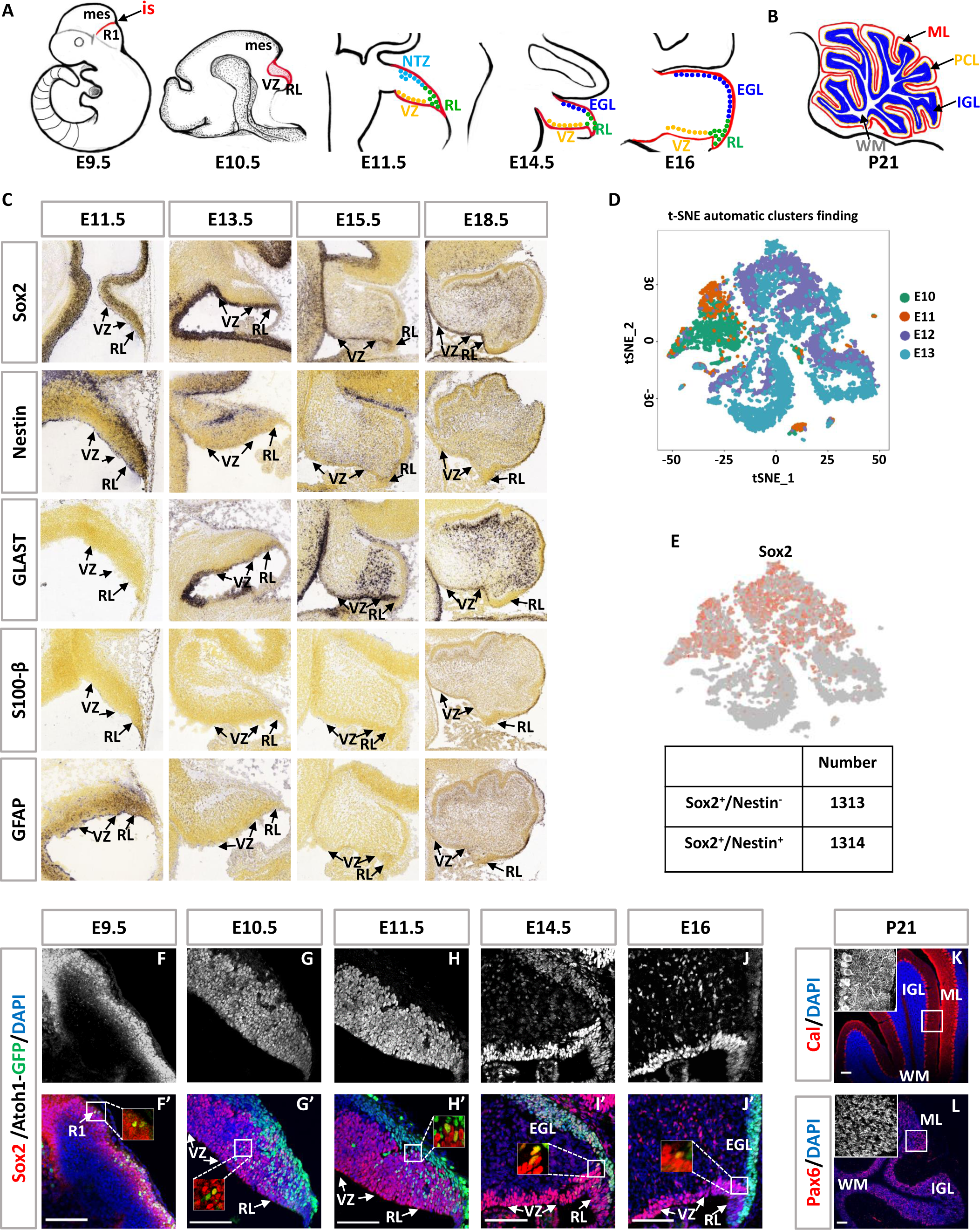
Sox2 is highly expressed both in the RL and VZ: (A-B) Cerebellar morphology at different stages during development from E9.5 to P21. R1, rhombomere1; is, isthmic organizer; mes, mesencephalon; RL, rhombic lip; VZ, ventricular zone; NTZ, nuclear transitory zone; EGL, external granule layer; IGL, internal granule layer; PCL, Purkinje cell layer; ML, molecular layer; WM, white matter. (C) In situ hybridization of the expression of Sox2, Nestin, GLAST, S100-β and GFAP (Data obtained from the Allen Brain Atlas). (D) t-distributed stochastic neighbor embedding (t-SNE) visualizations of cerebellar derived cell clusters (left) and developmental time points (right) at E10-E13. Each point represents one cell. Cells were color-coded according to time points. (E) t-SNE shows Sox2 expression and the cell number of Sox2^+^/Nestin^-^ cells and Sox2^+^/Nestin^+^ cells within Sox2^+^ cluster. Cells were color-coded according to genes expression. (F-J’) Immunostaining for Sox2 (gray for F-J, red for F’-J’) expression in a time series of mouse cerebellum during development. Squares in (F’-J’) represent Sox2^+^/Atoh1-GFP^+^ cells. Scalebars=100 μm. (K-L) Immunostaining for Calbindin (Cal, red) and Pax6 (red) expression in the cerebellum at P21. Scalebars=100 μm. Nuclei were stained with DAPI (blue).

**Figure S2.**
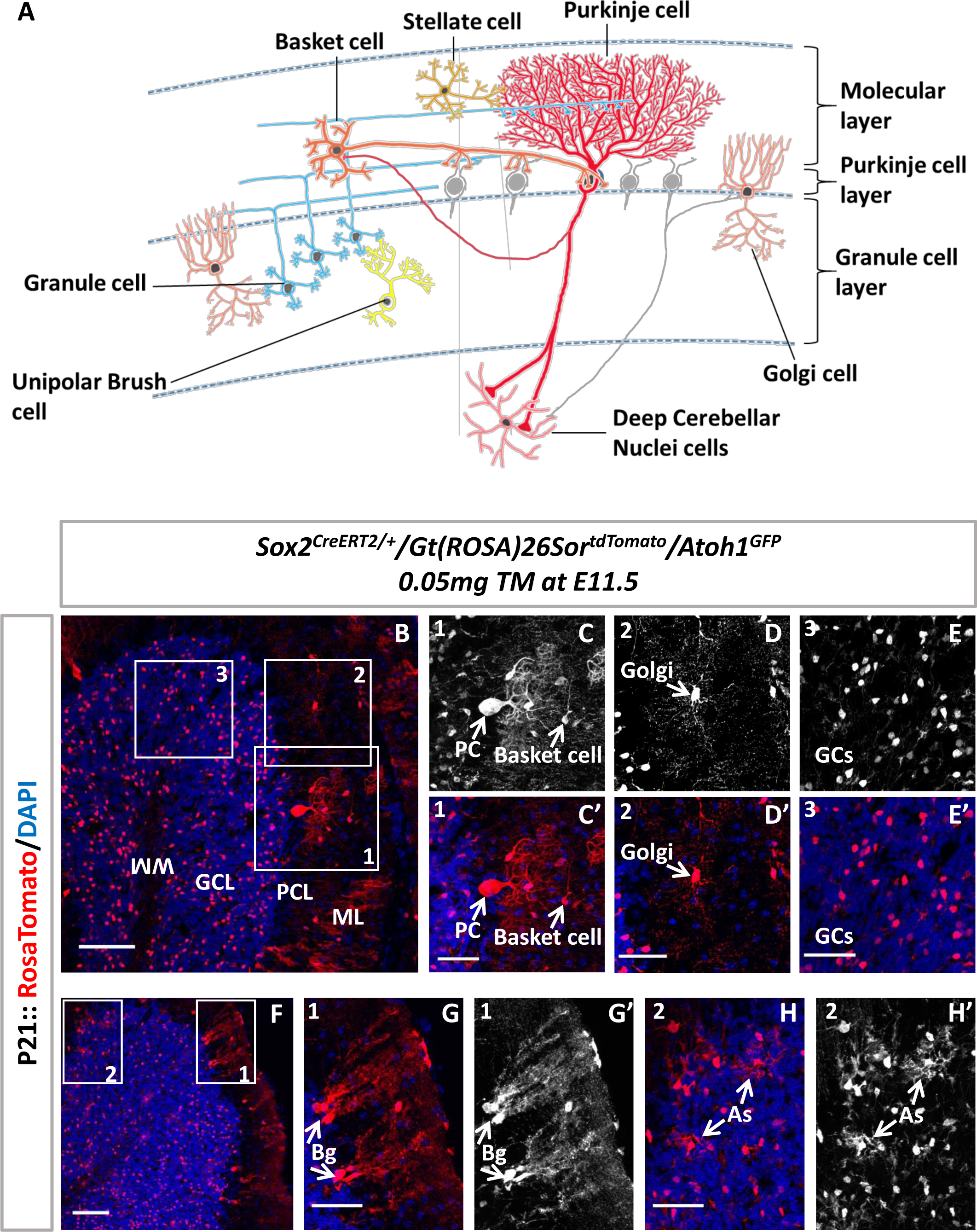
Cell diversity generation at P21: (A) Schematic of the morphology and position of glutamatergic and GABAergic neurons in the adult cerebellar cortex within a cerebellar folium. Purkinje cell is red, stellate cell is orange-yellow, basket cell is orange, Golgi cells are light orange, granule cells are blue, unipolar brush cell is yellow, deep cerebellar nuclei are orange-red. (B and F) Lineage tracing of Sox2^+^ progenitor clones (labeled by tdTomato, red) at P21. (C-E’ and G-H’) High magnification of the rectangular regions in (B and F) which show the Purkinje cell (PC, C and C’), Basket cell (C and C’), Golgi cell (D and D’), Granule cells (GCs, E and E’) Bergmann glia (Bg, G and G’) and Astroglia (As, H and H’). Nuclei were stained with DAPI (blue). Scalebars=100 µm and 50 µm.

**Figure S3.**
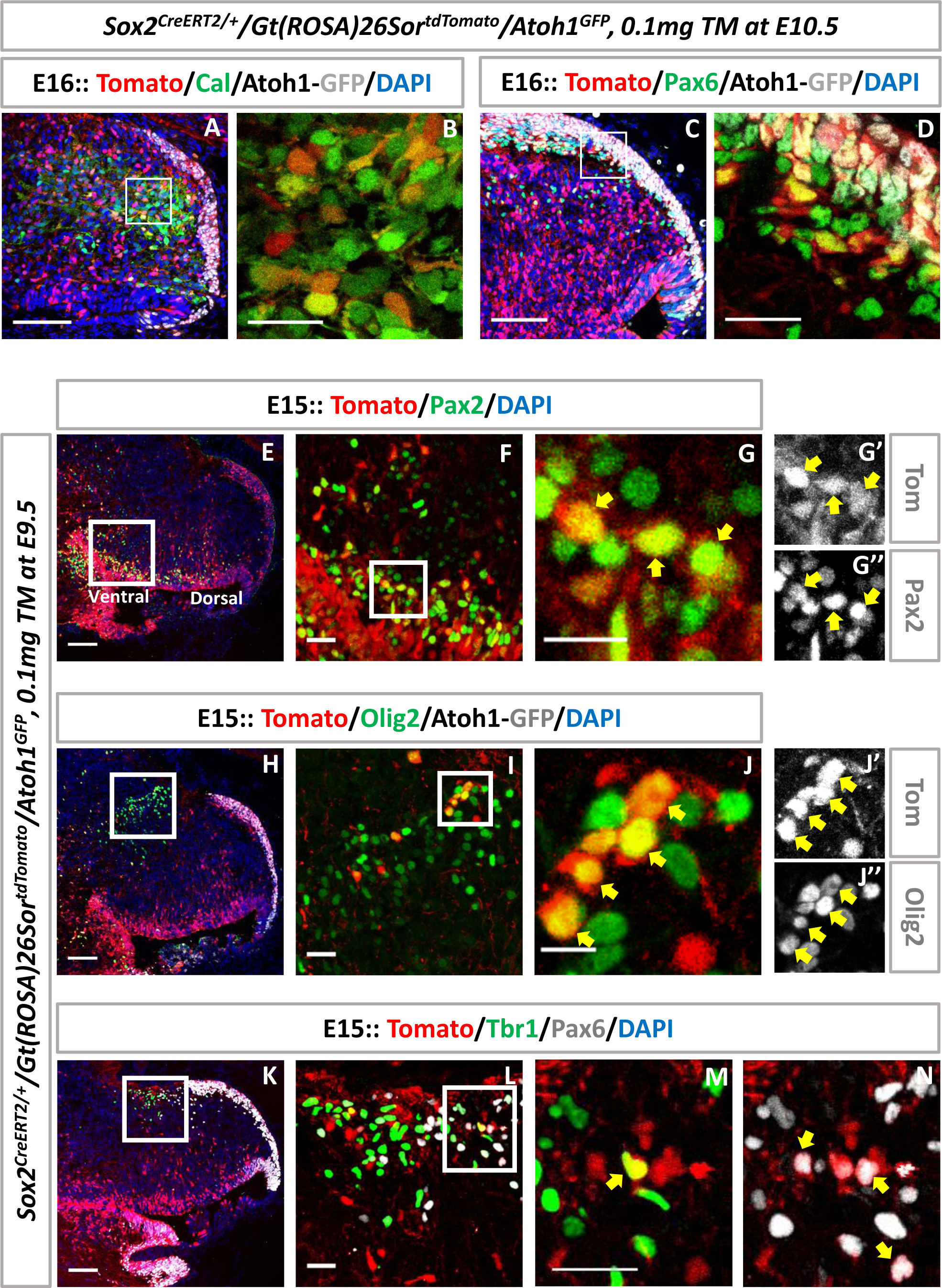
Sox2 progenitors give rise to the different cerebellar neuronal subtypes: (A and C) Co-localization of tdTomato (red) with Calbindin (green, A) or Pax6 (green, C) in mouse cerebellum at E16. Scalebars=100 µm. (B and D) High magnification of the rectangular regions in (A) and (C). Scalebars=25 µm. (E and F) Co-localization of Pax2 (green, a marker for INs) and tdTomato (red) in cerebellar primordium at E15. Scalebars=100 µm and 25 µm. (G-G’’) Higher magnification of the rectangular region in (F). Scalebars=10 µm. (H and I) Co-localization of Olig2 (green, a marker for glu-DCN) and tdTomato (red) at E15. Scalebars=100 µm and 25 µm. (J-J’’) Higher magnification of the rectangular region in (I). Scalebars=10 µm. (K and L) Triple immunolabeling with tdTomato (red), Pax6 (grey) and Tbr1 (green, a marker for glu-DCN) at E15. Scalebars=100 µm and 25 µm. (M and N) Higher magnification of the rectangular region in (L). Scalebars= 25 µm. Yellow arrows indicate double positive cells of Tomato^+^ with Pax2^+^ (G-G’’) or Olig2^+^ (J-J’’) or Pax6^+^ (N) or Tbr1^+^ (M). Nuclei were stained with DAPI (blue).

**Figure S4.**
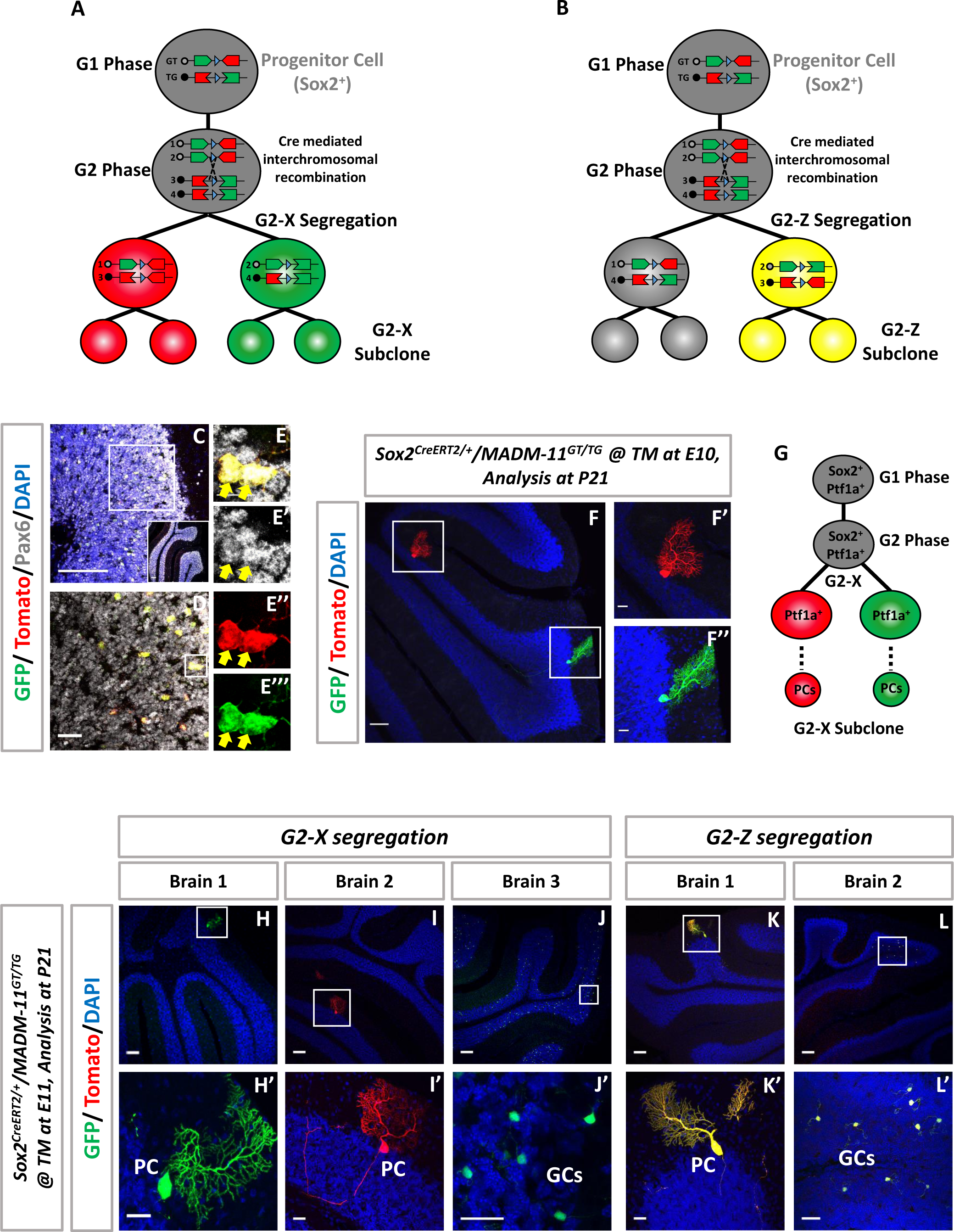
Single cell type of MADM clones in the mouse cerebellum: (A and B) Scheme of G2-X-MADM-clone (A) and G2-Z-MADM-clone (B) in MADM-11^GT/TG^/Sox2^CreER^ mice. (C and D) Immunostaining for Pax6, a marker for GCs, at P21. Scalebars=100 µm and 25 µm. (E-E’’’) Higher magnification of the rectangular region in (D). Scalebars=10 µm. (F) Single progenitor G2-X-MADM clone at P21. One green PC and one red PC were found in the same cerebellum. Scalebar=100 µm. (F’-F’’) Higher magnification of the rectangular regions in (F). Scalebars=25 µm. (G) Scheme of a single G2-X-MADM-clone generated fate restricted red (PCs) and green (PCs) lineages. (H-J) Single progenitor G2-X-MADM labeling cells in P21 cerebellum. Two PCs (green, H and red, I) and tens of green GCs (J) were found in different mouse cerebella. Scalebars=100 µm. (H’-J’) Higher magnification of the rectangular regions in (H-J). Scalebars=25 µm. (K and L) Single progenitor G2-Z-MADM labeling cells in P21 cerebellum. One yellow PC (K) and tens of yellow GCs (L) were found in different mouse cerebella. Scalebars=100 µm. (K’ and L’) Higher magnification of the rectangular regions in (K and L). Scalebars=25 µm. Nuclei were stained with DAPI (blue).

**Figure S5.**
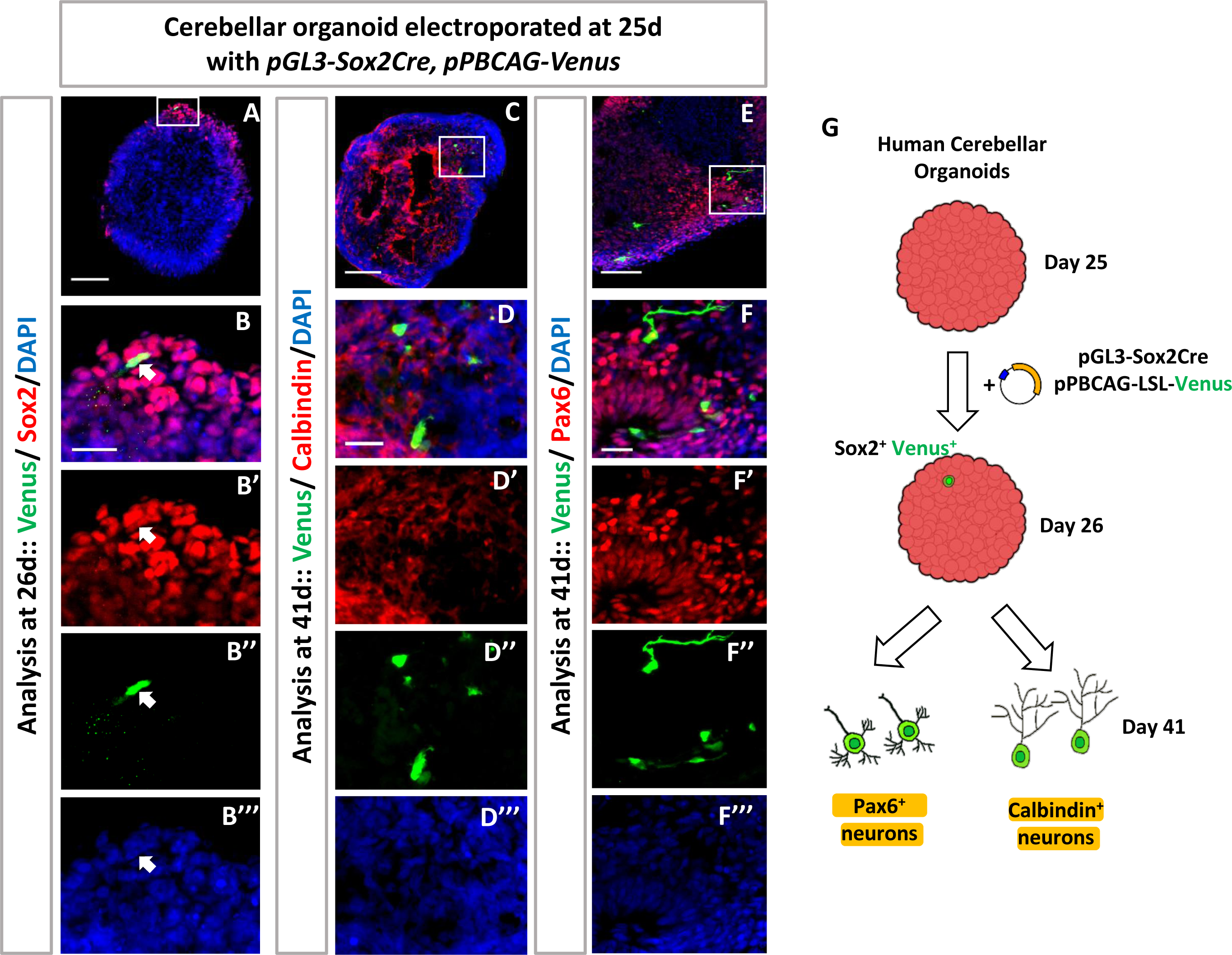
Sparse lineage tracing of ECPs in human cerebellar organoids. (A) Expression of Venus (green) and Sox2 (red) 1 day after electroporation. (B-B’’’) Higher magnification of the rectangular region in (A). Arrows indicate Sox2^+^/Venus^+^ double positive cells. (C and E) Immunolabeling of both PC marker (Calbindin, red) or GC marker (Pax6, red) co-localized with Venus (green) 16 days after electroporation. (D-D’’’ and F-F’’’) Higher magnification of the rectangular regions in (C and E). (G) Scheme of the experiments. Nuclei were stained with DAPI (blue). Scalebars=100 µm and 25 µm.

**Figure S6.**
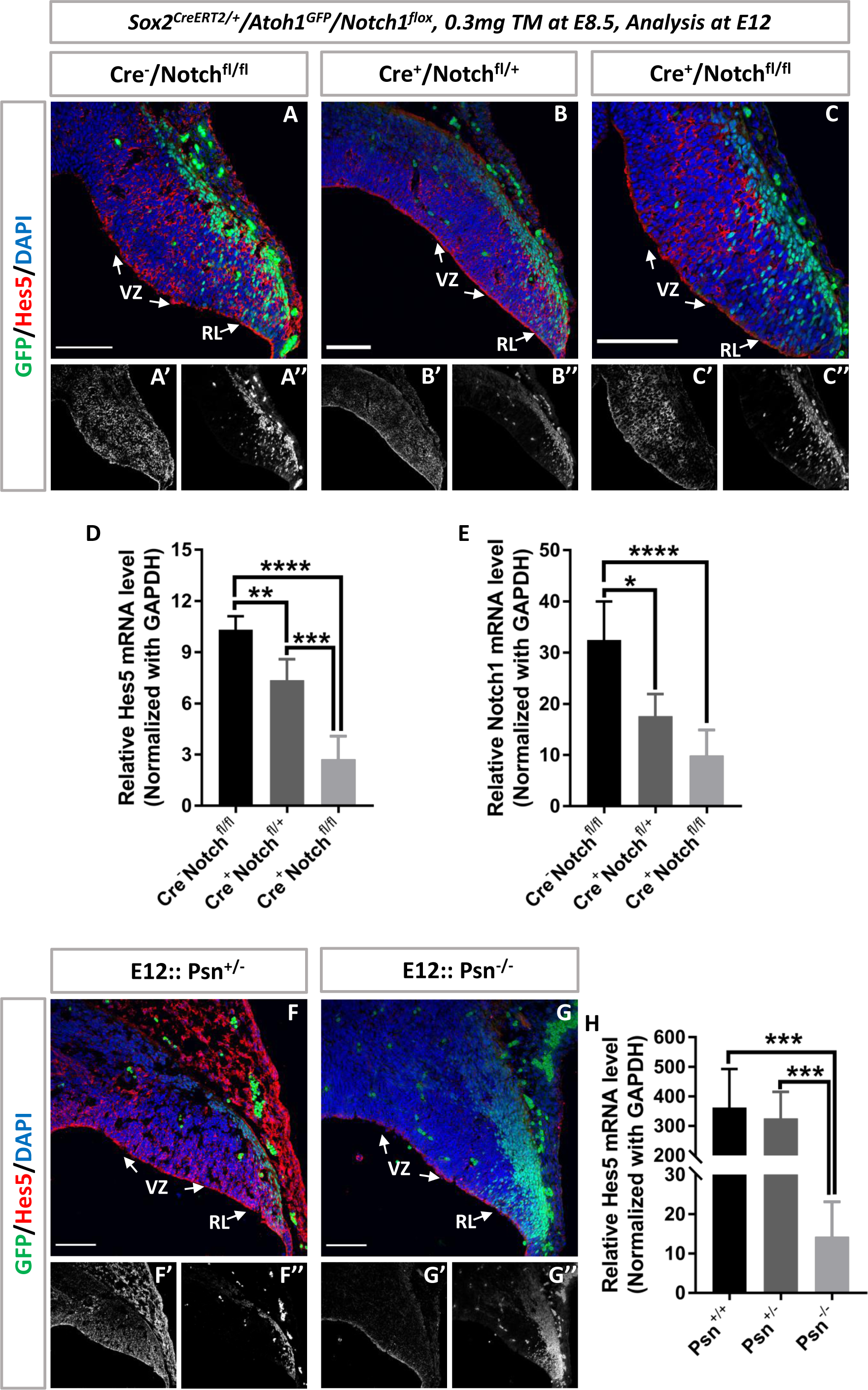
Notch loss of function decreases both Hes5 and Notch1 expression: (A-C’’) Immunostaining for Hes5 (red) at E12 in Cre^-^/Notch^fl/fl^ (A-A’’), Cre^+^/Notch^fl/+^ (B-B’’) and Cre^+^/Notch^fl/fl^ (C-C’’) cerebella. (D and E) Comparison of Hes5 and Notch1 mRNA levels in Cre^-^/Notch^fl/fl^ (n=6), Cre^+^/Notch^fl/+^ (n=6) and Cre^+^/Notch^fl/fl^ (n=6) mice. (F-G’’) Immunostaining for Hes5 (red) at E12 in Control (Psn^+/-^, F-F’’) and Presenilin1 (Psn) KO (Psn^-/-^, G-G’’) cerebella. (H) Comparison of Hes5 mRNA levels in Control (n=5 for Psn^+/+^ and n=6 for Psn^+/-^) and Presenilin1 (Psn) KO (n=5 for Psn^-/-^) cerebella. Nuclei were stained with DAPI (blue). Scalebars=100 µm. Data presented as mean ± SD. *p < 0.05, **p < 0.01, ***p < 0.001, ****p < 0.0001.

**Figure S7.**
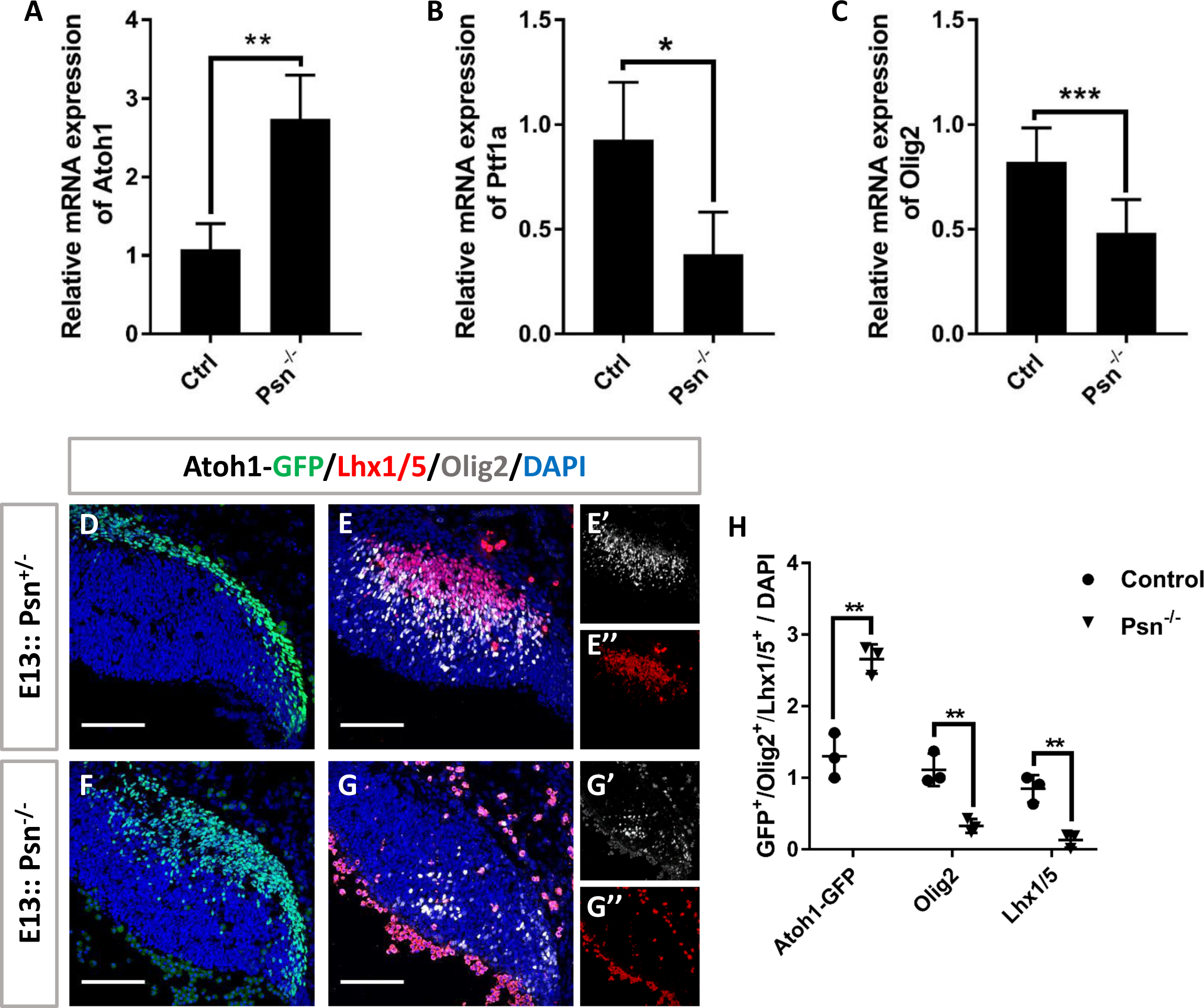
Notch loss of function increases RL progenitors at the expense of VZ progenitors. (A-C) Comparison of Atoh1 (n=4), Ptf1a (n=4) and Olig2 (n=10 for Control group and n=7 for Psn KO group) mRNA levels in Control and Psn KO cerebellum using RT-PCR at E13. (D) Immunostaining for Atoh1-GFP^+^ cells (green) at E13 in wild-type cerebellum. n=3. (E-E’’) Immunostaining for Lhx1/5^+^ cells (red) and Olig2^+^ (grey) at E13 in Control cerebellum. n=3. (F) Immunostaining for Atoh1-GFP^+^ cells (green) at E13 in Psn KO cerebellum. n=3. (G-G’’) Immunostaining for Lhx1/5^+^ cells (red) and Olig2^+^ (grey) at E13 in Psn KO cerebellum. n=3. (H) Percentage of Atoh1-GFP^+^ cells or Lhx1/5^+^ cells or Olig2^+^ cells compared with DAPI in Control and Psn KO cerebellum. Nuclei were stained with DAPI (blue). Scalebars=100 µm. Data presented as mean ± SD. *p < 0.05; **p < 0.01; ***p < 0.001.

**Figure S8.**
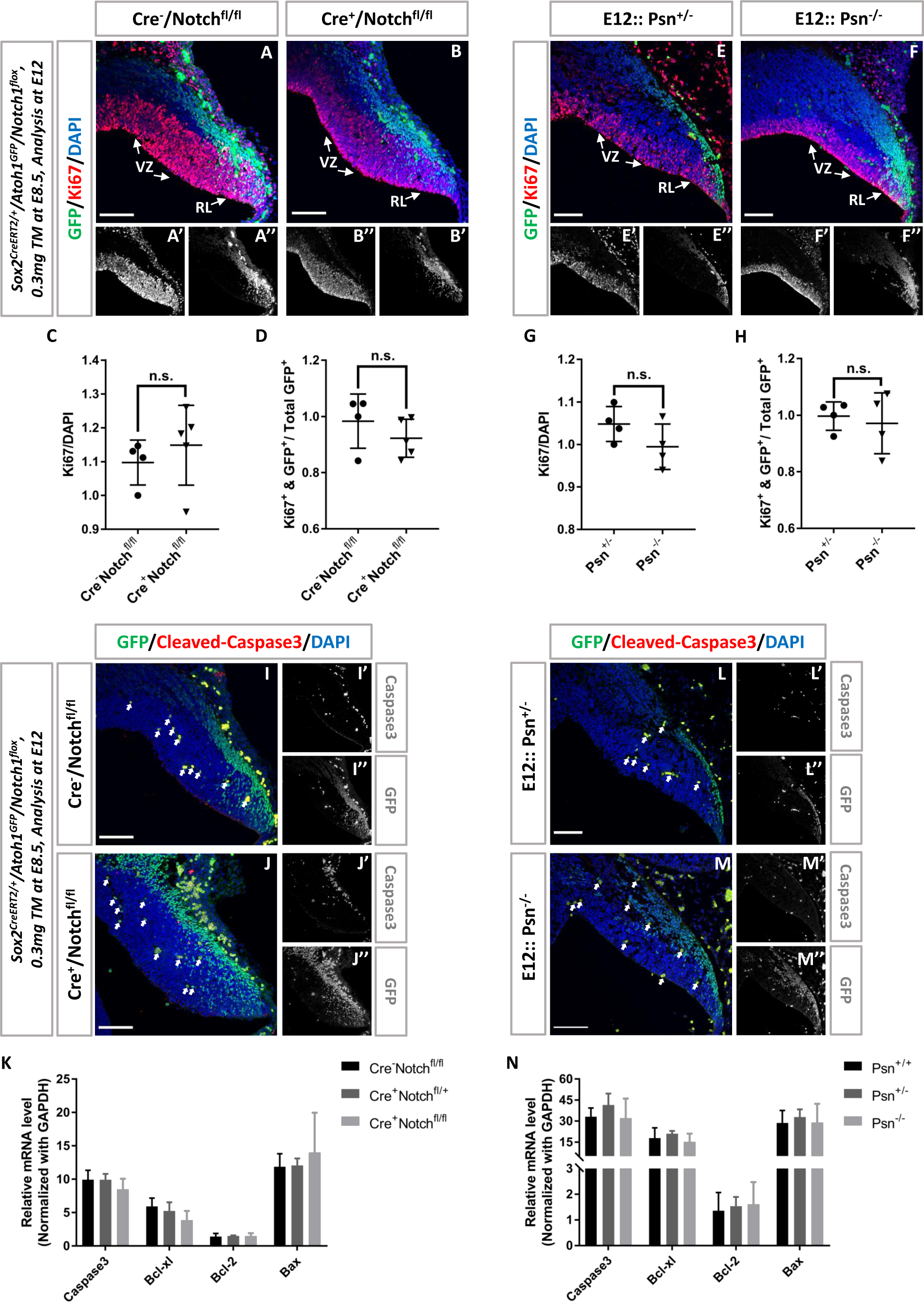
Notch loss of function does not affect cell proliferation and cell death in the cerebellar primordium. (A-B’’) Immunostaining for Ki67^+^ (red) at E12 in Cre^-^/Notch^fl/fl^ (n=4) and Cre^+^/Notch^fl/fl^ (n=5) mice. (C) Percentage of Ki67^+^ cells compared with DAPI. (D) Percentage of Ki67^+^/GFP^+^ double-positive cells within all Atoh1-GFP^+^ cells. (E-F’’) Immunostaining for Ki67^+^ (red) at E12 in Control (Psn^+/-^, n=4) and Presenilin1 (Psn) KO (Psn^-/-^, n=4) cerebella. (G) Percentage of Ki67^+^ cells compared with DAPI. (H) Percentage of Ki67^+^/GFP^+^ double-positive cells within all Atoh1-GFP^+^ cells. (I-J’’) Immunostaining for Cleaved-Caspase3^+^ (red) at E12 in Cre^-^/Notch^fl/fl^ and Cre^+^/Notch^fl/fl^ mice. (K) Comparison of cell death related genes mRNA level (Caspase3, Bcl-xl, Bcl-2 and Bax) in Cre^-^/Notch^fl/fl^ (n=6), Cre^+^/Notch^fl/+^ (n=3) and Cre^+^/Notch^fl/fl^ (n=6) mice using RT-PCR at E12. (L-M’’) Immunostaining for Cleaved-Caspase3^+^ (red) at E12 in Control (Psn^+/-^) and Presenilin1 (Psn) KO (Psn^-/-^) cerebella. (N) Comparison of cell death related genes mRNA level (Caspase3, Bcl-xl, Bcl-2 and Bax) in Psn^+/+^ (n=5), Psn^+/-^ (n=5) and Psn^-/-^ (n=6) mice using RT-PCR at E12. Nuclei were stained with DAPI (blue). Scalebars=100 µm. Data presented as mean ± SD. n.s. > 0.05.

**Figure S9.**
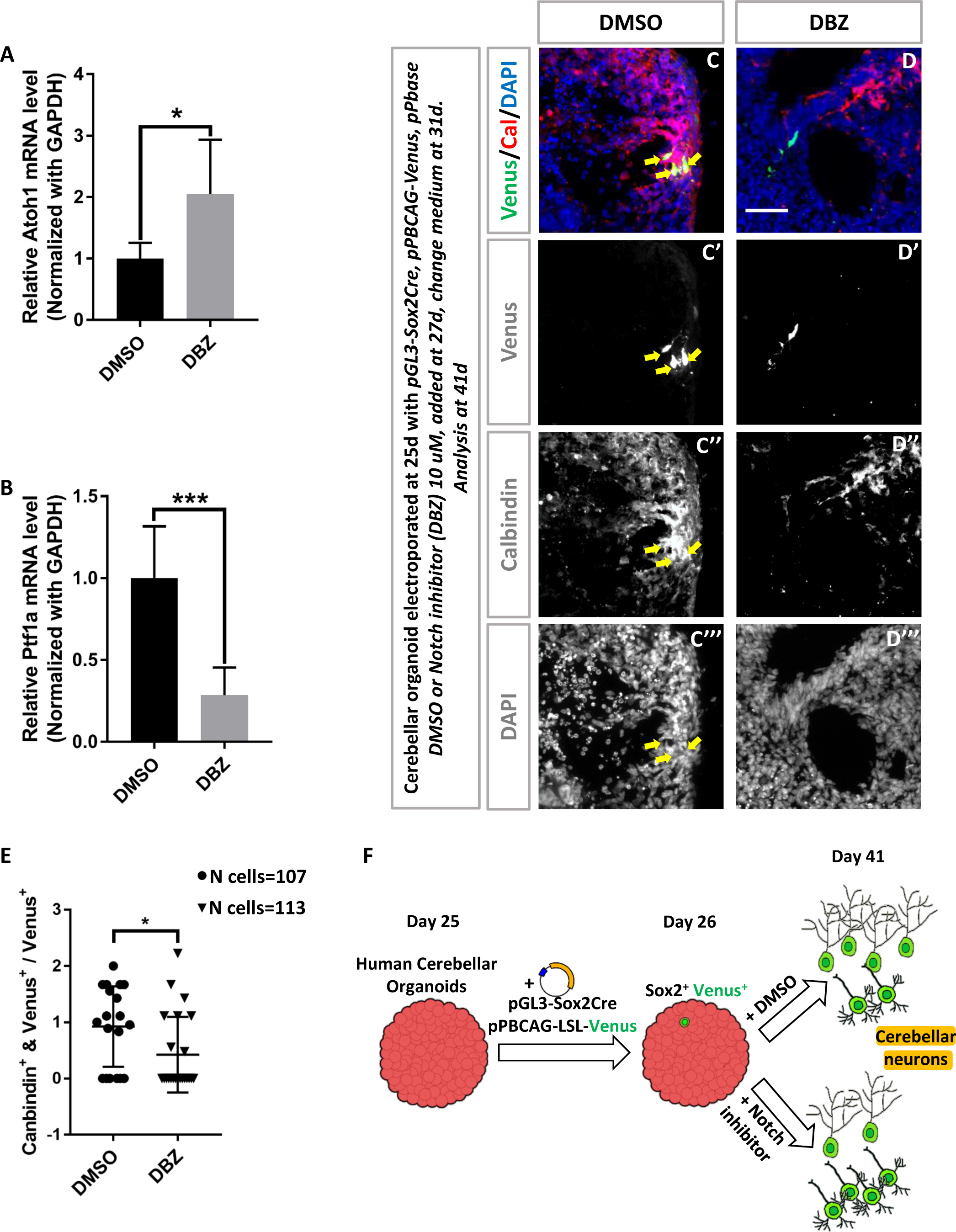
Notch inhibition increases glutamatergic progenitors at the expense of GABAergic progenitors in human cerebellar organoids: (A and B) Comparison of Atoh1 (n=6) and Ptf1a (n=6) mRNA level in DMSO treated control groups and Notch inhibitor (DBZ) treated groups in human cerebellar organoids using RT-PCR. (C-D’’’) Immunolabeling of PC marker (Calbindin, red) co-localized with Venus (green) 16 days after electroporation in DMSO treatment group (C-C’’’, n=19) and in DBZ (Notch inhibitor, 10µM) treatment group (D-D’’’, n=23). (E) Percentage of Calbindin^+^ and Venus^+^ double positive cells within all Venus^+^ cells in DMSO versus DBZ treatment groups. (F) Scheme of the experiments. Arrows indicate Calbindin^+^/Venus^+^ double positive cells. Nuclei were stained with DAPI (blue). Scalebars=50 µm. Data presented as mean ± SD. *p < 0.05.

**Figure S10.**
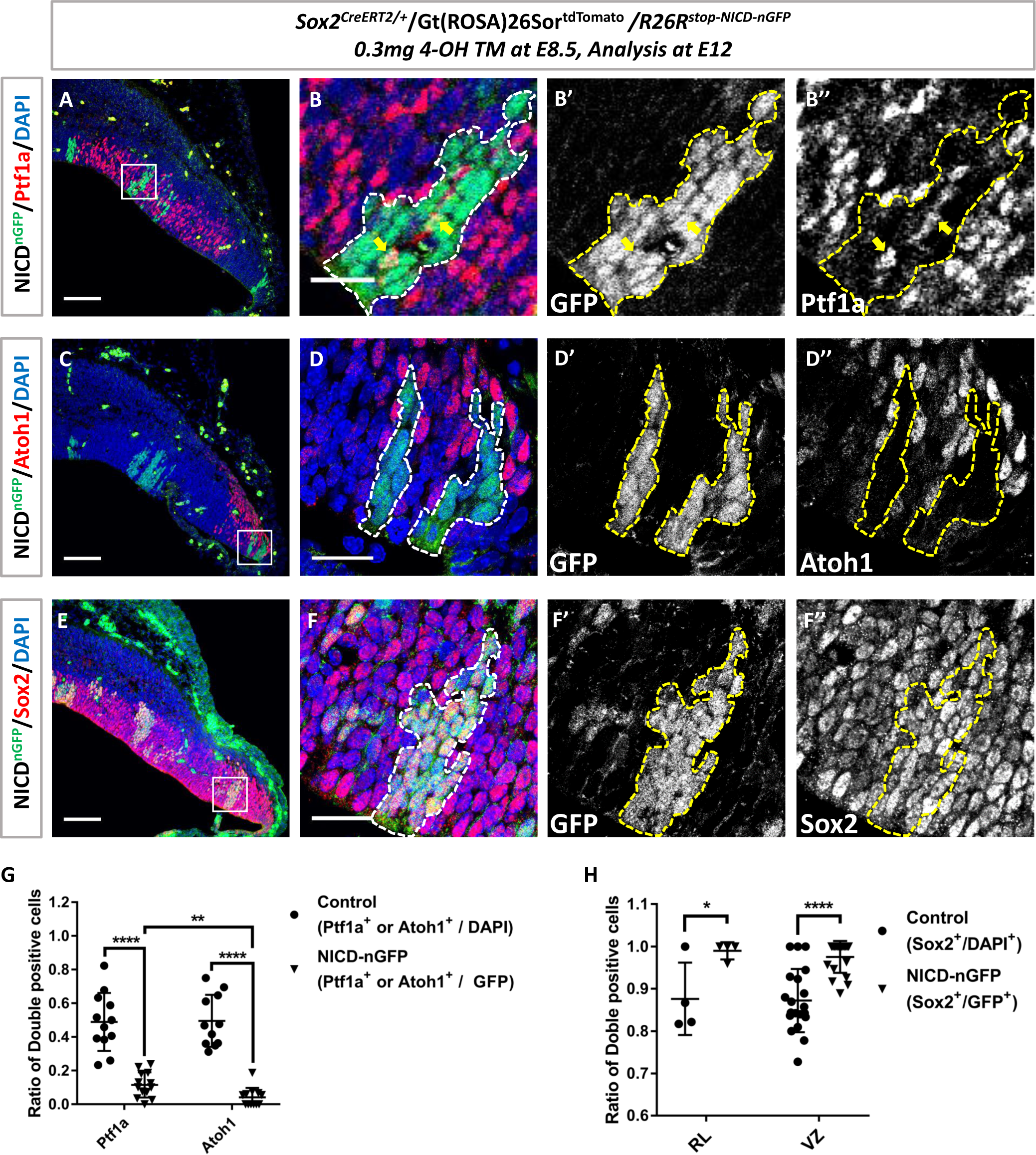
Notch GOF in ECPs inhibits their differentiation compared with adjacent wild-type cells: (A) Immunostaining for Ptf1a^+^ (red) in GFP^+^ Notch GOF clones (green) at E12. n=4 embryos, totally 12 clones in the VZ. (B-B’’) Higher magnification of the rectangular region in (A). Arrows indicate NICD-GFP^+^/Ptf1a^+^double positive cells. (C) Immunostaining for Atoh1^+^ (red) in GFP^+^ Notch GOF clones (green) at E12. n=4 embryos, totally 11 clones in the RL. (D-D’’) Higher magnification of the rectangular region in (C). (E) Immunostaining for Sox2^+^ (red) in GFP^+^ Notch GOF clones (green) at E12. n=4 embryos, totally 4 clones in the RL and 19 clones in the VZ. (F-F’’) Higher magnification of the rectangular region in (E). The ratio of Ptf1a^+^ cells or Atoh1^+^ cells to total cells in GFP^+^ Notch GOF clones (Ptf1a^+^ or Atoh1^+^/ GFP^+^) compared to the ratio of Ptf1a^+^ cells or Atoh1^+^ cells to an equivalent number of adjacent control cells (Ptf1a^+^ or Atoh1^+^/ DAPI^+^) in the VZ or RL, respectively. (H) The ratio of Sox2^+^ cells to total cells in GFP^+^ Notch GOF clones (Sox2^+^/GFP^+^) compared with the ratio of Sox2^+^ cells to an equivalent number of adjacent control cells (Sox2^+^ / DAPI^+^) both in the RL and VZ. Nuclei were stained with DAPI (blue). Scalebars=100 µm and 25 µm. Data presented as mean± SD. *p < 0.05; **p < 0.01; ****p < 0.0001.

**Figure S11.**
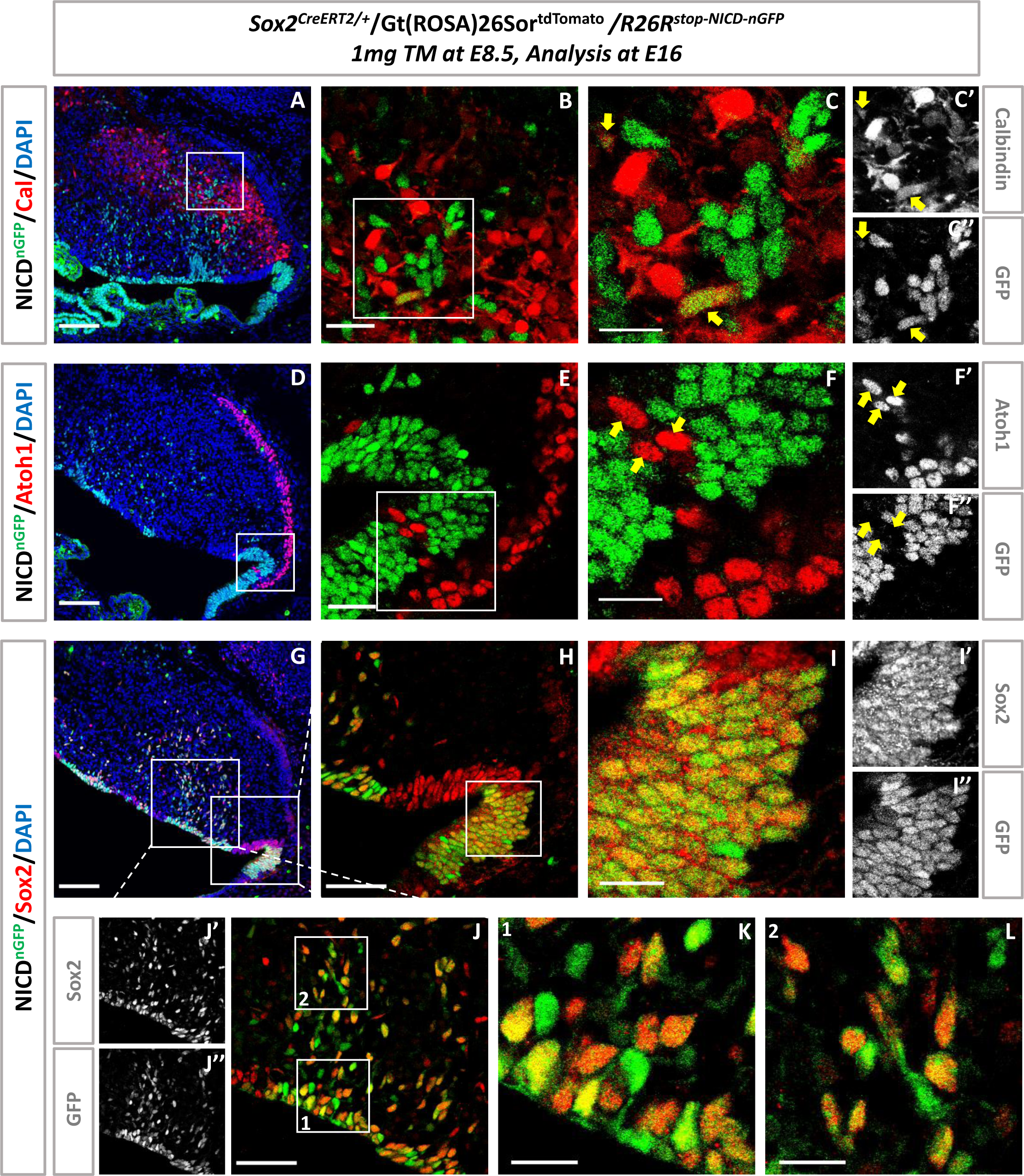
Notch GOF in ECPs shows lasting inhibition of their differentiation: (A and D) Immunostaining for Calbindin (A) and Atoh1 (D) at E16, Scalebars=100 µm. (B and E) Higher magnification of the rectangular region in (A and D), Scalebars=25 µm. (C-C’’ and F-F’’) Higher magnification of the rectangular region in (B and E), Scalebars=15 µm. Arrows indicate NICD-GFP^+^/Ptf1a^+^double positive cells (C-C’’) and NICD-GFP^-^ /Atoh1^+^ cells (F-F’’). (G) Immunostaining for Sox2 at E16, Scalebars=100 µm. (H) Higher magnification of the rectangular region in (G, RL region). Scalebars=25 µm. (I-I’’) Higher magnification of the rectangular region in (H). Scalebars=15 µm. (J-J’’) Higher magnification of the rectangular region in (G, VZ region). Scalebars=25 µm. (K and L) Higher magnification of the rectangular region in (J). Scalebars=15 µm. Nuclei were stained with DAPI (blue).

**Table S1.** scRNAseq analysis of the mean level of Notch signaling related genes at E12 and E13.

|  | Ptf1a only |  | Sox2 only |  | Atoh1 only |  |
| --- | --- | --- | --- | --- | --- | --- |
|  | E12 | E13 | E12 | E13 | E12 | E13 |
| Number of cells | 479 | 370 | 898 | 975 | 1375 | 665 |
| Mean (Hes5) | 2.0106 | 1.72 | 1.9952 | 2.1 | 1.1337 | 0.5 |
| Mean (Hes1) | 0.4288 | 0.6121 | 0.6756 | 1.1664 | 0.3307 | 0.2595 |
| Mean (Notch1) | 0.3358 | 0.3026 | 0.3659 | 0.3639 | 0.2625 | 0.0847 |
| Mean (Dll1) | 0.6101 | 0.8748 | 0.3115 | 0.2302 | 0.5419 | 0.3557 |
| Mean (Dll3) | 0.211 | 0.1502 | 0.1325 | 0.1115 | 0.8623 | 0.771 |
| Mean (Dll4) | 0.0012 | 0.0049 | 0.0022 | 0.005 | 0.0042 | 0.0172 |
| Mean (Jag1) | 0.0218 | 0.0241 | 0.0264 | 0.039 | 0.0328 | 0.0336 |
| Mean (Jag2) | 0 | 0 | 0.0007 | 0 | 0.0011 | 0 |

**Table S2.**
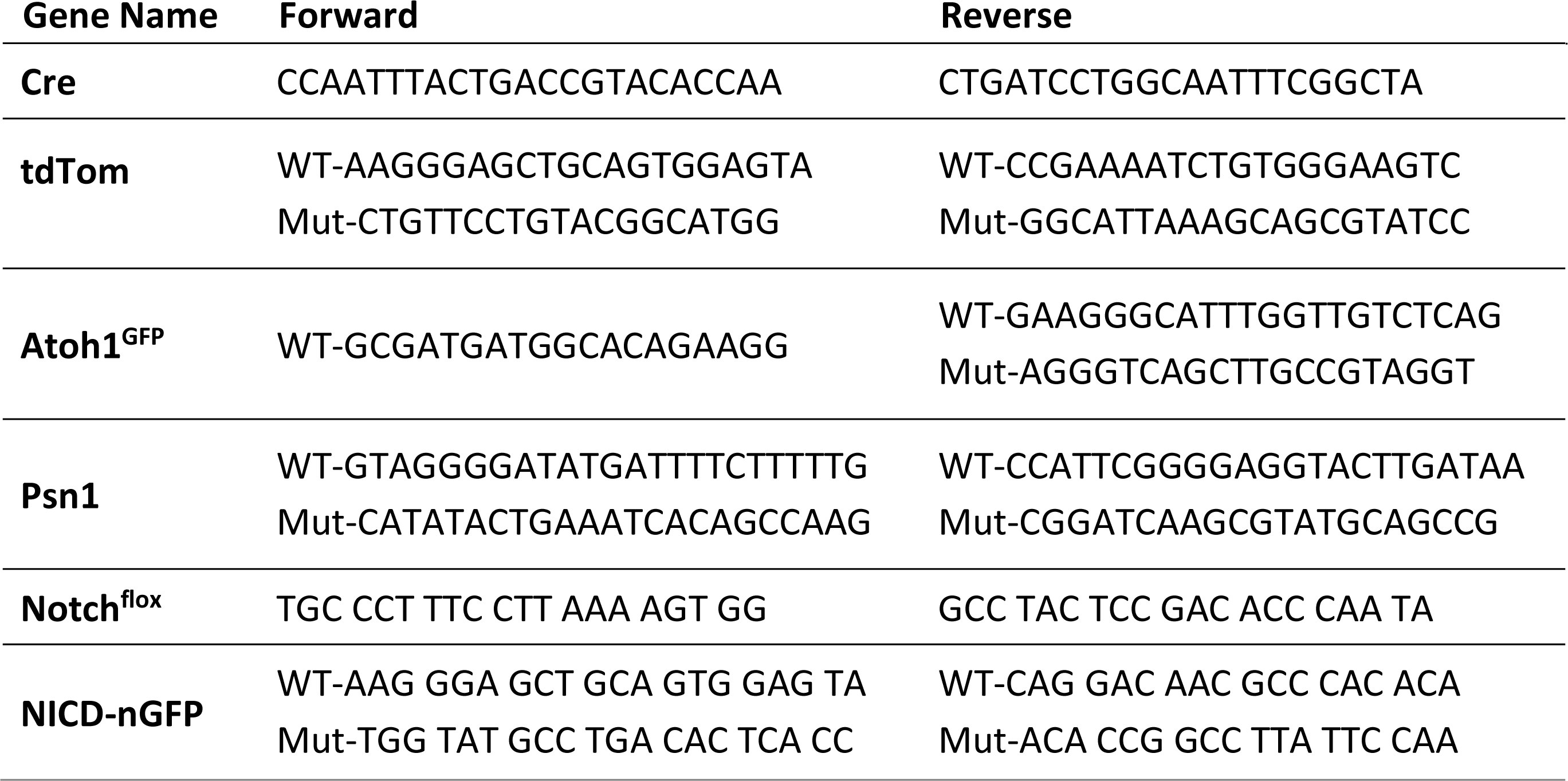
Primers for mouse genotyping.

**Table S3.**
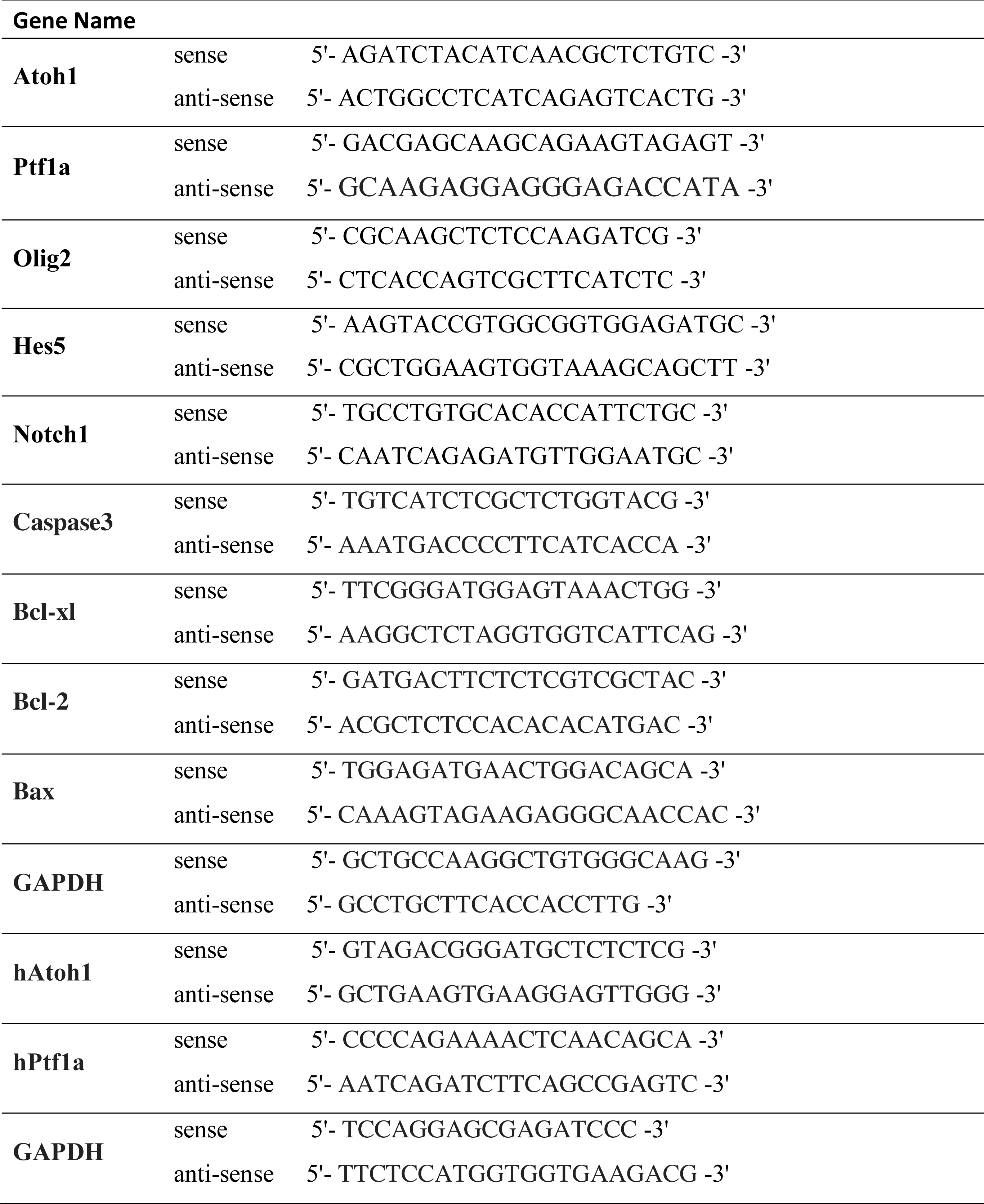
Primers for RT-PCR.

